# Ultra High-Plex Spatial Proteogenomic Investigation of Giant Cell Glioblastoma Multiforme Immune Infiltrates Reveals Distinct Protein and RNA Expression Profiles

**DOI:** 10.1101/2022.10.04.510833

**Authors:** Shilah A. Bonnett, Alyssa Rosenbloom, Giang Ong, Mark Conner, Aric Rininger, Daniel Newhouse, Felicia New, Chi Phan, Saskia Ilcisin, Hiromi Sato, John Lyssand, Gary Geiss, Joseph M. Beechem

## Abstract

A deeper understanding of complex biological processes, including tumor development and immune response, requires ultra high-plex, spatial interrogation of multiple “omes”. Here we present the development and implementation of a novel spatial proteogenomic (SPG) assay on the GeoMx® Digital Spatial Profiler platform with NGS readout that enables ultra high-plex digital quantitation of proteins (> 100-plex) and RNA (whole transcriptome, > 18,000-plex) from a single FFPE sample. This study highlighted the high concordance, *R* > 0.85, and <11% change in sensitivity between SPG assay and the single analyte –assays on various cell lines and tissues from human and mouse. Furthermore, we demonstrate that the SPG assay was reproducible across multiple users. When used in conjunction with advanced cellular neighborhood segmentation, distinct immune or tumor RNA and protein targets were spatially resolved within individual cell subpopulations in human colorectal cancer and non-small cell lung cancer. We used the SPG assay to interrogate 23 different glioblastoma multiforme samples across 4 pathologies. The study revealed distinct clustering of both RNA and protein based on pathology and anatomic location. The in-depth investigation of giant cell glioblastoma multiforme revealed distinct protein and RNA expression profiles compared to that of the more common glioblastoma multiforme. More importantly, the use of spatial proteogenomics allowed simultaneous interrogation of critical protein post-translational modifications alongside whole transcriptomic profiles within the same distinct cellular neighborhoods.

## Introduction

The advancement of spatially resolved, multiplex proteomic and transcriptomic technologies has revolutionized and redefined the approaches to complex biological questions pertaining to tissue heterogeneity, tumor microenvironments, cellular interactions, cellular diversity, and therapeutic response (1). These spatial technologies, including the GeoMx® Digital Spatial Profiler (DSP), can yield spatially resolved proteomic and transcriptomic datasets from formalin-fixed paraffin-embedded (FFPE) or fresh frozen (FF) samples. Most of these approaches are specific towards generating either proteomic or transcriptomic datasets. Multiple studies have demonstrated a poor correlation between RNA expression and protein abundance in samples when each analyte is profiled, with the most egregious of cases often owing to target- or tissue-specific post-transcriptional regulation (2). Despite our current understanding of the variety of mechanisms surrounding transcriptional and translational regulation, target turnover, post-translational protein modifications, and protein activity, RNA is still the primary analyte of choice in highly multiplexed studies. A workflow that accurately measures RNA and protein simultaneously within a single sample and spatial context is critical to a fuller understanding of the global state of the cell.

Previously, to understand proteogenomic and transcriptomic relationships, researchers would acquire individual analyte-specific datasets, often employing different technologies, and computationally integrate them using various multiomic approaches (3–6) (Fig. 1A). While this workflow provides a deeper understanding of the biological system under study, the potentially confounding variables associated with variation stemming from employing multiple technologies, section-to-section variability, or precisely matching regions of interest (ROIs) across multiple slides must be taken into consideration when analyzing and interpreting the data. To control or eliminate these potentially confounding technical variables, multimodal omics, defined as the simultaneous co-detection of multiple analytes (‘omes’) in a single sample, serves as an advantageous alternative approach (7–11). In the emerging spatial biology field, there has been a growing interest in the development of novel multimodal omic protocols to detect RNA transcript levels and protein abundance within a single sample while maintaining the spatial context within a tissue. These novel multimodal omic datasets of protein and DNA or RNA have been termed “spatial proteogenomics”. However, until recently, spatial proteogenomic protocols with ultrahigh-plex, simultaneous co-detection of analytes have not been available (8–21). Many of the protocols combine high-plex RNA with up to 4 fluorescent antibodies utilized as morphology markers.

**Figure 1:**
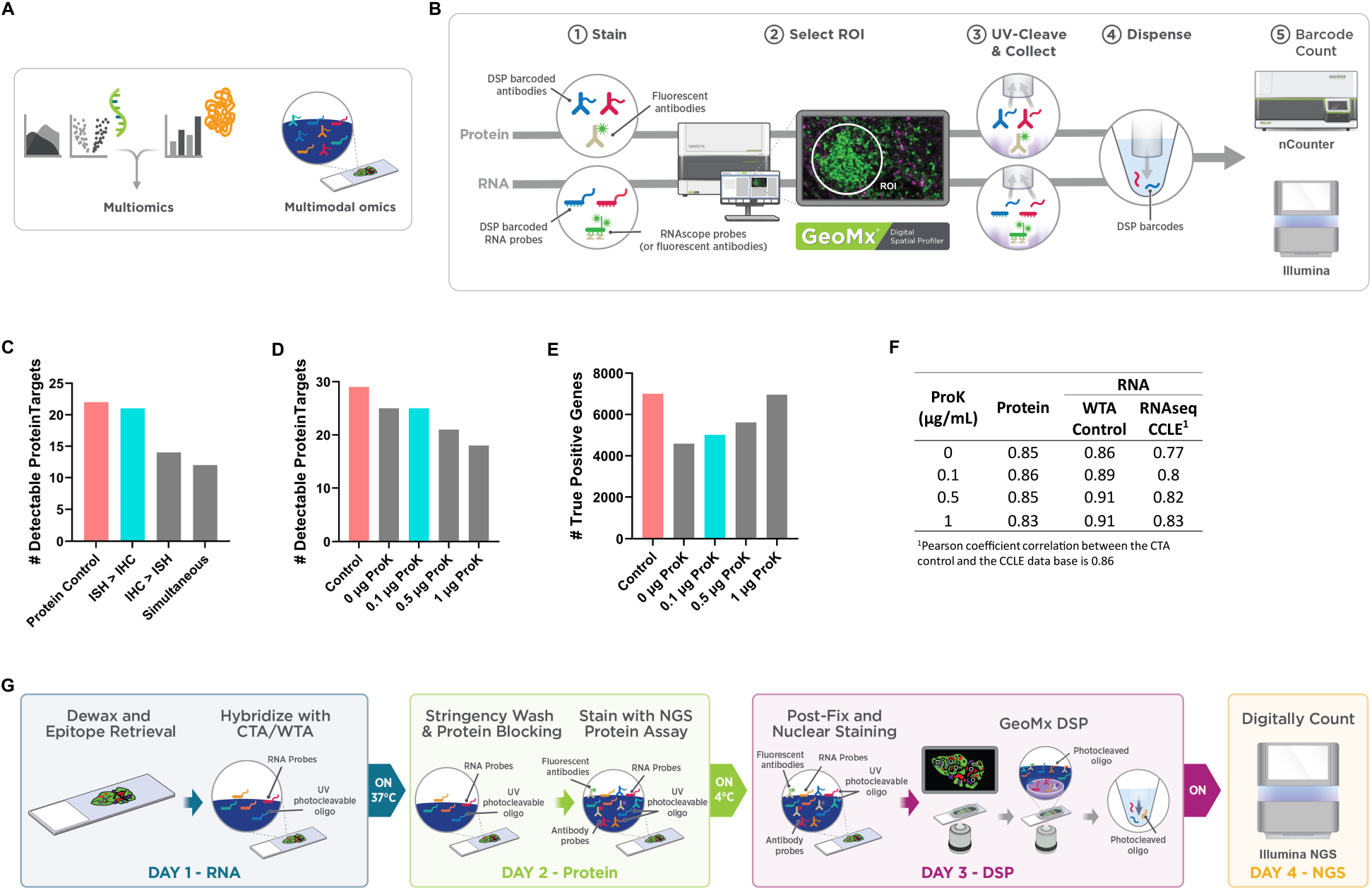
Technical development of the spatial proteogenomic assay. (**A**) Current proteogenomic approaches are multiomic which entails the integration of individual-omic datasets and multimodal which involves the simultaneous, co-detection of multiple ‘omes’ in a single sample. (**B**) Commercially available GeoMx Assays currently enable high-plex, spatially resolved protein and RNA targets on individual tissue sections with nCounter or NGS quantitative readout. (**C**) Assessment of staining order on the number of protein targets above detection threshold. FFPE cell line, A431CA, was stained with GeoMx Protein assays (59-plex) for nCounter readout and mock RNA probe (Buffer R only). Assessment of varying proteinase K on the performance of the spatial proteogenomic assay. A 45-cell pellet array (CPA) was stained with 59-plex GeoMx NGS Protein modules (59-plex) and GeoMx WTA under proteogenomic and standard assay conditions. Plots represents the number of targets above detection threshold for (**D**) protein (SNR > 3) and (**E**) true positives detectable WTA targets. (**F**) Pearson’s correlation on log_2_ transformed SNR data between the proteogenomic assay and the single analyte controls along with the CCLE RNAseq database. Circular ROIs of 200 μm diameter were selected for detailed molecular profiling with the GeoMx DSP. The signal was averaged across replicate AOIs and the SNR was calculated. (**G**) GeoMx Spatial Proteogenomic workflow enables multimodal omic profiling on a single slide.

The GeoMx DSP enables spatially resolved, high-plex digital quantitation of proteins (> 100-plex) and RNA (whole transcriptome, > 18,000-plex) from human and mouse tissues (22–25). Previous work has detailed proteomic or transcriptomic single-analyte workflows using the GeoMx DSP technology with unique affinity reagents (antibodies for protein or *in situ* hybridization (ISH) probes for RNA) coupled to photocleavable oligonucleotide barcodes (22,23,25) (Fig. 1B). Tissue samples are incubated with one of these affinity reagents, and oligonucleotide barcodes are precisely liberated from an area of interest (AOI) with UV light exposure. The released barcodes are then collected for quantification with next-generation sequencing (NGS). While the assays were developed for single-analyte analysis, there is no fundamental technical barrier to concurrently profile protein and RNA targets on a single sample. However, the divergent sample preparation protocols for RNA and protein would hinder the concurrent profiling of both analytes. Here we introduce a novel GeoMx DSP spatial proteogenomic workflow that allows for the simultaneous co-detection and quantification of RNA transcripts and protein abundance in a defined cell population within individual FFPE mouse and human tissues, using oligonucleotide-barcoded affinity reagents and the GeoMx DSP platform and Illumina NGS readout. We demonstrate the performance of the assay on various cell lines and tissues and highlight its use in a biological context on various glioblastoma multiforme samples including giant cell glioblastoma (gcGBM).

## Results

### Spatial Proteogenomic Assay Development

Multiomic profiling of RNA and Protein generally requires two serial tissue sections (Fig. 1B). Serial sections often include similar, but not identical, cell populations and thus do not provide the optimal measurement of the relationship between gene expression and protein abundance. Therefore, we set out to develop a novel high-plex spatial multi-modal assay on the GeoMx DSP platform with NGS readout, which allows for the simultaneous profiling of high-plex RNA transcripts and proteins from a defined cell population within individual AOI on a single FFPE tissue section. We named this assay GeoMx Spatial Proteogenomic Assay or SPG for short.

For the individual GeoMx NGS Protein assay and GeoMx NGS RNA assay, sample preparation protocols are nearly identical to standard immunohistochemistry (IHC) or *in situ* hybridization (ISH) methodologies, respectively. The GeoMx NGS Protein assay uses a single antigen retrieval process of a slightly acidic heat-induced epitope retrieval (HIER) buffer (pH 6.0) under high pressure. The GeoMx NGS RNA assay uses a two-step, tissue-dependent epitope retrieval process with a basic HIER buffer (pH 9.0) followed by a proteolytic-induced epitope retrieval (PIER) step. Given two analytes that required distinct and disparate antigen retrieval conditions, we first optimized sample treatment conditions compatible with both analytes: staining, epitope retrieval, and Proteinase K digestion.

### Staining Strategy

Sample preparation for ISH normally involves harsh conditions, such as high salt concentrations and prolonged exposure to formamide at elevated temperatures, all of which may disrupt the antigen-antibody complex and thus reduce protein detection in FFPE tissue samples. To investigate the impact of ISH conditions on protein antigen detection, we first evaluated two sequential staining strategies: ISH staining followed by IHC staining (ISH > IHC) and in the reverse order (IHC > ISH). After implementing both staining steps, slides were subsequently processed on the GeoMx platform. We hypothesized ISH > IHC would result in optimal protein detection when compared to the reverse order. When IHC staining was performed first, followed by ISH, we found a slightly lower correlation (*R* = 0.86) and a 36% decrease in sensitivity when compared to single-analyte protein control (Fig. 1C and Supplementary Fig. S1). In contrast, carrying out ISH first, followed by IHC staining, had only a minor impact on protein correlation (*R* = 0.95) and sensitivity (5% decrease).

We also evaluated a simultaneous strategy, concurrently staining with both antibodies (for IHC) and RNA probes (for ISH) under conditions where formamide was reduced 5-fold. For this experiment we evaluated the impact of the simultaneous staining conditions on protein detection in absence of RNA probes. While formamide allows for the hybridization to occur at lower temperatures and reduces the non-specific binding of RNA probes, it can disrupt antibody-antigen interactions, and thus the quality of antibody-based protein detection (8,31). With the simultaneous staining strategy, we observed a 45% decrease in protein target detection, indicating disruption of antibody-antigen binding even in the reduced formamide concentration (Fig. 1C and Supplementary Fig. S1). Therefore, we concluded that the optimal strategy for dual detection of RNA and protein targets was sequential staining with ISH followed by IHC.

### Impact of Epitope Retrieval Conditions

Next, we sought to identify the optimal epitope retrieval conditions for the SPG assay. The standard GeoMx RNA and Protein assays are designed to perform at opposing HIER conditions, basic and slightly acidic conditions, respectively, whereas the SPG assay workflow calls for a single epitope retrieval condition. To optimize the HIER conditions specifically for the SPG assay, eleven FFPE cell-pellet array (CPA) sections were pretreated under basic or slightly acidic HIER conditions followed by PIER with 1 μg/mL of Proteinase K (ProK). The CPA sections provides a uniform set of cells with known RNA expression levels as defined by the CCLE RNAseq dataset (32). Pretreated sections were then stained in a sequential fashion with the GeoMx Human Cancer Transcriptome Assay (GeoMx CTA; ~1,800 protein-coding genes) followed by a set of modular GeoMx Human NGS Protein panels (59-plex) (Supplementary Table S3). For each cell line, the signal was averaged across biological replicate ROIs and the signal-to-noise ratio (SNR) was calculated for protein and RNA targets. The performance of the SPG assay was compared to the single-analyte assay control slides.

When the SPG assay data on FFPE cell lines were compared to the single analyte RNA assay controls, we observed a strong correlation (*R* > 0.95) regardless of slightly acidic or basic HIER pretreatment conditions (Supplementary Fig. S2A and S2B). Additionally, HIER pretreatment conditions had little impact on the correlation between the SPG assay data and the Cancer Cell Line Encyclopedia (CCLE) RNAseq data (*R* > 0.84) (32) (Supplementary Fig. S2C). From the CCLE RNAseq dataset, we identified a true set of expressed genes (TPM > 1) that was used to calculate the true positive rate (TPR; sensitivity) and false positive rate (FPR, specificity) with respect to assessed RNA targets. The FPR of the SPG assay under basic HIER conditions was < 10% (Supplementary Fig. S2D). In contrast, for the assay under slightly acidic HIER conditions, the FPR increased to 30%. The high FPR associated with slightly acidic HIER conditions is consistent with the previous observation: an increase in non-specific hybridization when epitope retrieval was performed under slightly acidic conditions (25).

The SPG assay was then compared to the single-analyte protein assay from FFPE cell lines. The pretreatment of samples with slightly acidic HIER demonstrated a higher correlation *(R* = 0.86) to the protein assay control than with the basic HIER treatment (*R* = 0.77) (Supplementary Fig. S3A and S3B, respectively). Thus, the evaluation of epitope retrieval methods indicated optimal detection of protein under slightly acidic HIER conditions and the optimal detection of RNA under basic conditions, which is consistent with the standard GeoMx single-analyte workflows. While detection sensitivity for protein decreased when slides were pretreated under basic conditions followed by ProK, nonspecific binding for RNA increased under slightly acidic conditions. To move forward, we chose to maintain RNA detection specificity and refine protein sensitivity by titrating ProK concentrations under basic HIER conditions.

### Impact of Proteinase K concentrations

We noted the relatively high concentration of ProK (1 μg/mL) used in the PIER step drove protein target detection loss due to antigen digestion (Supplementary Fig. S3C). Thus, we assessed the effects of ProK concentration under basic HIER on protein and RNA target detection in the SPG assay. To evaluate the effects of various concentrations of ProK, 45 FFPE CPA sections were stained with the GeoMx Human Whole Transcriptome Atlas (GeoMx Human WTA; >18,000 protein-coding genes) probe set and a 59-plex set of GeoMx Human NGS Protein panels. FFPE cell lines were assessed under basic HIER conditions followed by PIER with varying concentrations of ProK. When comparing the single-analyte protein control with the SPG assay, the SNR correlations between these two assays remained relatively strong (*R* = 0.83-0.86, Fig. 1F and Supplementary Fig. S4A-D) and FPR remained < 10%, regardless of ProK concentrations (Supplementary Fig. S4E). However, the levels of protein target detection significantly decreased (> 37%) at ProK concentrations > 1 μg/mL (Fig. 1D). At 0.1 μg/mL ProK, the number of detected protein targets by the SPG assay was comparable (~13% decrease in detection) to the single-analyte protein control.

For RNA targets, the correlation between the SPG assay and the RNA single-analyte control was higher when the samples were treated with increased ProK concentrations as expected (*R* = 0.86 for 0 μg/mL ProK, *R* = 0.89-0.91 for 0.1-1.0 μg/mL ProK, Fig. 1F and Supplementary Fig. S5A-D). A similar trend was observed when we compared the SPG assay data of RNA targets to the CCLE RNAseq data (*R* = 0.77 for 0 μg/mL ProK, *R* = 0.8-0.83 for 0.1-1.0 μg/mL ProK, Fig. 1F and Supplementary Fig. S5E). Using the CCLE RNAseq dataset, we identified a true set of expressed genes (TPM > 1) that was used to calculate the number of true positives. In our analysis, the number of true positives increased with higher concentrations of ProK (Fig. 1E and Supplementary Fig. S5F). These results demonstrate the critical balance between ProK proteolytic digestion and optimal RNA detection. Even at the lowest ProK concentrations, digestion of critical protein epitopes was detectable. Particularly, the detection of low abundance protein and RNA targets was the most affected by ProK.

### GeoMx Spatial Proteogenomic Workflow

After optimizing the staining strategy for the SPG assay, including the epitope retrieval conditions and ProK concentrations on sensitivity and specificity for protein and RNA detection, we established the optimal GeoMx SPG workflow. This workflow consists of a sequential staining strategy of ISH followed by IHC under basic (pH 9.0) HIER conditions with the ProK digestion (PIER) step at a low concentration (0.1 μg/mL). The optimized GeoMx SPG workflow requires a total of 4 days to complete from the slide preparation step to the data acquisition and analysis, which is described below. The GeoMx SPG workflow is compatible only with NGS readout including custom NGS probes and not compatible with nCounter® readout (Fig. 1G).

The overall SPG workflow for FFPE samples is as follows:

- **Day 1**: A two-step epitope retrieval process involving HIER under basic (pH 9.0) conditions followed by a PIER step using 0.1 μg/mL ProK. Samples are then incubated with the GeoMx WTA or GeoMx CTA RNA probe cocktails overnight for hybridization at 37°C.
- **Day 2**: Samples are washed under stringent conditions in the presence of 50% formamide and subsequently treated with a blocking solution to prevent nonspecific antibody binding. After blocking, samples are stained with one or more GeoMx NGS Protein panel(s) overnight at 4°C. Fluorophore-conjugated primary antibodies may be added at this step to visualize tissue morphology.
- **Day 3**: After fixing with 4% paraformaldehyde and staining with a nuclear marker (Syto13), samples are processed on the GeoMx DSP and then sequenced on the Illumina NextSeq2000 or Illumina NovaSeq6000 as noted in Materials and Methods.
- **Day 4**: Process data with the GeoMx NGS Pipeline as described in Materials and Methods.

### Profiling of Cell Lines using the Optimized GeoMx Spatial Proteogenomic Workflow

Using the optimized GeoMx SPG assay workflow, we profiled FFPE CPA sections using the GeoMx Human WTA and the 15-modular GeoMx Human NGS Protein panel simultaneously (147-plex) (Supplementary Table S3). The GeoMx Human Protein Assays with NGS readout enables to profile of up to 144 protein targets (plus additional three IgG controls) simultaneously. As protein and RNA controls, CPAs were independently stained with the standard single-analyte GeoMx Human NGS Protein panel (147-plex) or with the standard single-analyte GeoMx Human WTA, respectively. Protein targets were considered detected if the SNR was ≥ 3. To ensure we were well above the noise when making comparisons for the RNA targets, we implemented an SNR cutoff of ≥ 4.

During the technical development of the SPG assay, we used a combination of 59-plex Human NGS protein panels (including IgG controls). Once the SPG assay was developed, we increased the number of panels to 15, composed of 147 proteins (147-plex, including IgG controls). By profiling FFPE of 45 different cell lines, the correlation of these two assays was tested. The SNRs of the SPG assay data using the 147-plex protein panel demonstrated a strong correlation (*R* = 0.92) between proteins that overlap with the original 59-plex protein panel (Fig. 2A). This finding suggests that more than doubling the number of proteins measured had little impact on assay performance.

**Figure 2:**
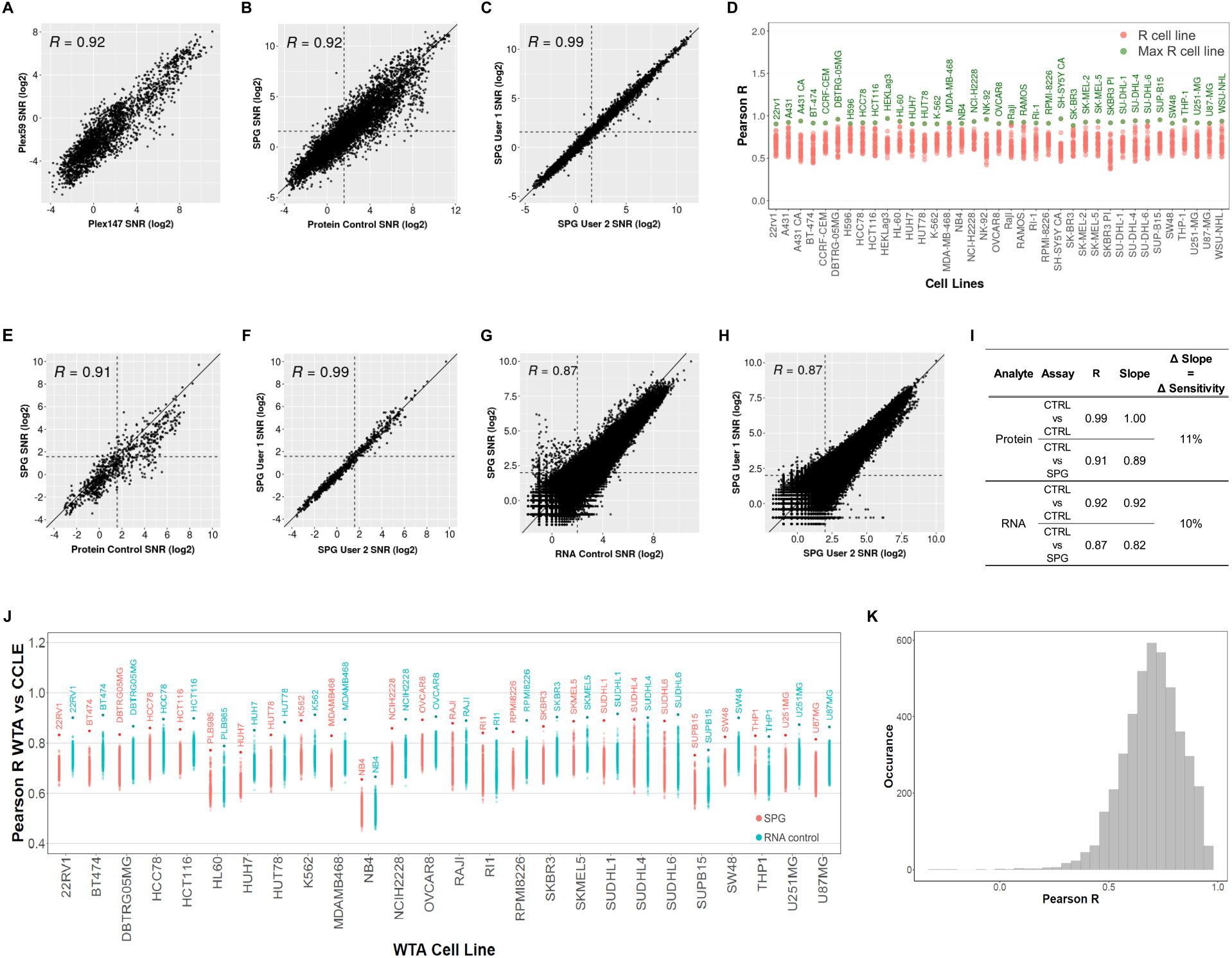
Assessment of spatial proteogenomic data quality versus the respective RNA and Protein control data. Assay correlation with respect to the protein analyte comparing (**A**) 147-plex and 59-plex protein panel, (**B**) protein control and proteogenomic workflows and (**C**) user-to-user and instrument-to-instrument reproducibility. (**D**) Cell line to cell line comparison of Protein Control to proteogenomic protein data. For protein targets with SNR > 3, the Pearson’s *R* was calculated between each cell line from the Protein Control slide against all the cell lines in the spatial proteogenomic slide. The max R cell line between the SPG and Protein control is labeled and highlighted green. Assay correlation of 17 phospho-specific antibodies between (**E**) protein control and proteogenomic workflows and (**F**) user-to-user and instrument-to-instrument reproducibility. Assay correlation with respect to the RNA analyte comparing (**G**) RNA control and proteogenomic workflows and (**H**) user-to-user and instrument-to-instrument reproducibility. (**I**) Summary of Pearson’s R, slope of linear regression and change in sensitivity between workflows. The change in sensitivity corresponds to the average change in regression line slope between the SPG and the single analyte control assay. (**J**) Cell line to cell line comparison of WTA Control and proteogenomic WTA data to the entire CCLE RNA-Seq dataset. For all overlapping targets between the CCLE and WTA data, the Pearson’s *R* in the Protein Control and spatial proteogenomic WTA data were calculated against all cell lines in the CCLE RNA-Seq. Cell line labels in the plot correspond to SPG or GeoMx WTA cell lines with the highest R correlation to the CCLE data. (**K**) Target-to-target comparison of WTA control to proteogenomic WTA data. For each RNA target with SNR > 4 in 15% of samples, the Pearson’s *R* was calculated between WTA control log2 SNR transformed data and the respective proteogenomic WTA log2 SNR transformed data. Histogram shows the distribution of Pearson’s *R*.

Using the 147-plex Human NGS protein panel, we compared the performance of the SPG assay to the protein control assay. There was a strong correlation (*R* = 0.92) between the two workflows (Fig. 2B). Furthermore, both the single analyte protein and SPG assay was reproducible across multiple users and instruments, where a very strong correlation (*R* = 0.99) was observed (Fig. 2C, Supplementary Fig. S6A). A pairwise correlation analysis was performed across 37 cell lines and all detectable targets. In the cell line to cell line comparison, the Pearson’s *R* was calculated between each cell line from the SPG assay against all cell lines in the single analyte control. A dot plot was generated showing the Pearson’s *R* distribution for each cell line. There was a high correlation between the same cell lines in these two assays (Fig. 2D). Additionally, we observed a high correlation between protein targets regardless of assay type in the target-to-target comparison across cell lines (Supplementary Fig. S7). To preserve the phosphorylation state of phosphorylated proteins, cell lines SKBR3, A431, and SH-SY5Y, making up the 45 CPAs, had been treated with a phosphatase inhibitor prior to paraformaldehyde fixation. The inhibitor treatment of cell lines partially drives a high correlation among phospho-specific antibodies as observed in the target-to-target heatmap (Supplementary Fig. S7).

For cellular activities, protein phosphorylation plays a critical role in the regulation of various cellular processes including cell signaling, gene expression, and cell growth and differentiation. Aberrant phosphorylation events are associated with several diseases including cancer, neurodegenerative disorders, and metabolic disorders. While transcriptomics enables the comprehensive profiling of cell and tissue specific gene expression, it is unable to decipher the phosphorylation state of key proteins. To study phosphorylation-related diseases and biological activities, using phospho-specific antibodies is one way to capture the phosphorylation state of the proteins. In the GeoMx Human NGS Protein panels, approximately ~12% of the validated antibodies are designed to detect and quantify phospho-specific proteins. The performance of the phospho-specific protein detection from FFPE cell lines under SPG assay conditions was similar to the single-analyte protein control (*R* = 0.91) (Fig. 2E, Supplementary Fig. S8). Furthermore, the performance of the phospho-specific detection with the SPG assay was reproducible across multiple users and instruments (*R* = 0.99) (Fig. 2F, Supplementary Fig. S8).

We evaluated the change in sensitivity, as denoted by the average change in regression line slope, of the SPG assay compared to the single analyte protein control (Fig. 2I). Using homogenous cell pellets, minimized the confounding variables associated with section-to-section variability and precisely matching ROIs across multiple tissue samples. In our analysis, we observed a 11% decrease in sensitivity for the SPG assay.

In addition to evaluating the detection of protein targets, we examined the data quality of the RNA targets detected with the SPG assay in comparison to the RNA control and the CCLE RNAseq dataset. A strong correlation (*R* = 0.87) was observed when comparing the single-analyte RNA control with the SPG assay (Fig. 2G). As with the protein analyte, there was a high correlation (*R* = 0.87) between multiple users and instruments for both the SPG and the single analyte RNA assay (Fig. 2H, Supplementary Fig. S6B). Furthermore, we observed a 10% decrease in sensitivity for SPG assay compared to the single analyte control (Fig. 2I). For all overlapping targets between the GeoMx Human WTA and the CCLE RNAseq dataset, each cell line in the proteogenomic assay and the RNA control were correlated to every cell line in the CCLE dataset (1,012 cell lines). For both the SPG and RNA control assays, we observed the highest correlation between matched cell lines to the CCLE RNAseq dataset (Fig. 2J). The target-to-target comparison of single-analyte WTA control to SPG WTA data indicated a high correlation between RNA targets between assay types (Fig. 2K).

### GeoMx Spatial Proteogenomics Assay in Human Tissue

Having validated the SPG assay on idealized cell pellet samples, we sought to evaluate the assay on human colorectal cancer (CRC) samples. We profiled matched immune (CD45^+^) and tumor (PanCK^+^) regions in serial FFPE tissue sections stained with the GeoMx Human WTA and/or the 147-plex modular GeoMx Human NGS Protein panels. For these experiments, geometric ROIs (100 μm diameter circles) were collected for each morphology specific regions (Fig. 3A and Supplementary Fig. S9A). In CRC samples, we observed a high correlation (*R* > 0.81) across ROIs between the SPG assay as compared to the single-analyte protein (Fig. 3B) and RNA assays (Fig. 3C). The correlations were higher in cell lines; however, this is to be expected given confounding variables associated with section-to-section variability or precisely matching ROIs across multiple slides. We then performed a correlation analysis on all CD45-enriched or PanCK-enriched ROIs between the SPG assay and the single-analyte protein or RNA assays. A dot plot was generated showing the Pearson’s *R* distribution for each ROI. In this analysis, we observed a high correlation between immune ROIs and between tumor ROIs from the SPG and single analyte controls (Supplementary Fig. S10A and B).

**Figure 3:**
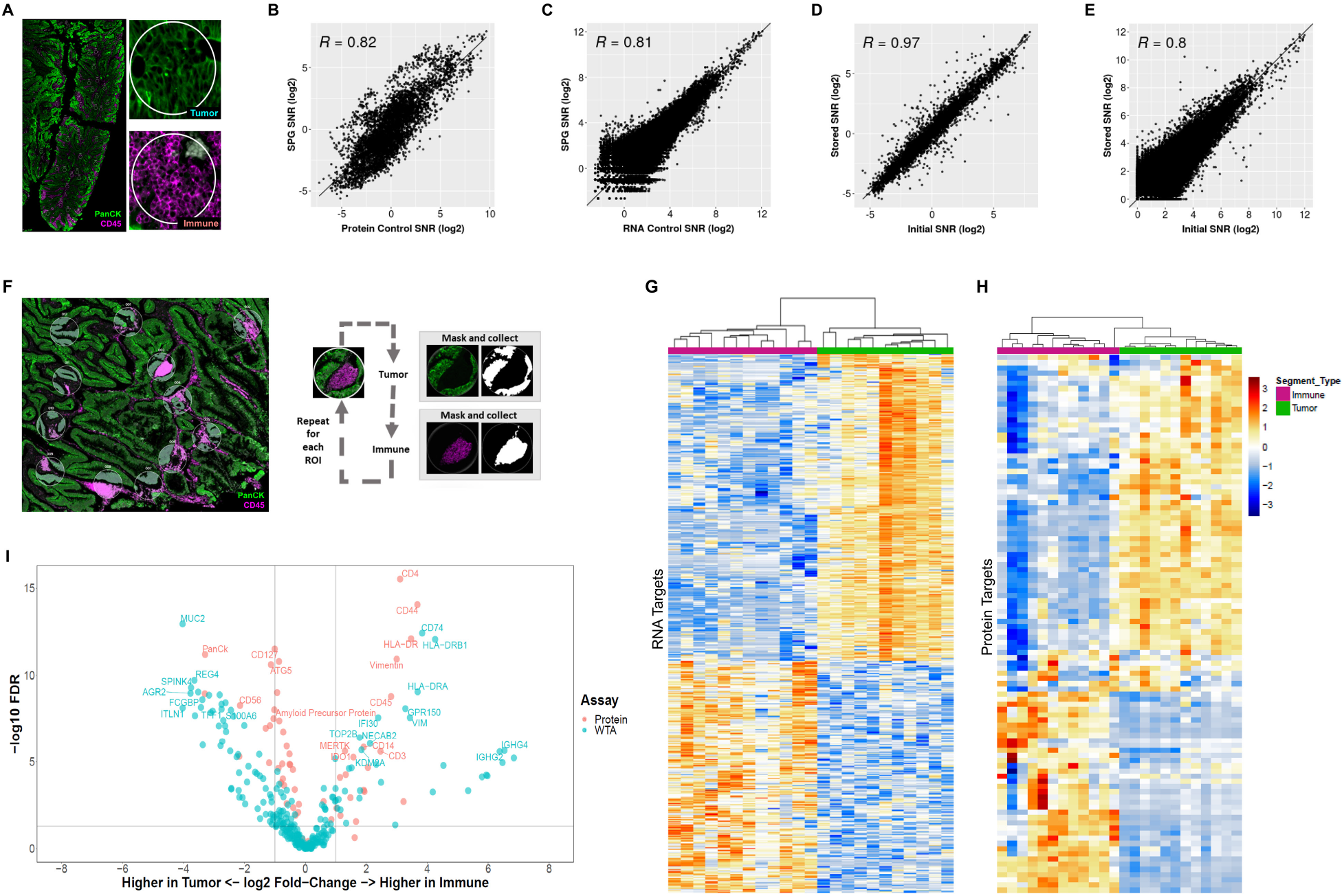
Assessment of spatial proteogenomic performance on human tissue. FFPE colorectal cancer (CRC) sections were stained with the GeoMx NGS Human Protein modules (147-plex), WTA, and antibodies against PanCK (Tumor) and CD45 (Immune). (**A**) Representative image of CRC sample used in the assessment of spatial proteogenomic data quality versus the respective RNA and Protein control data. Two ROIs showing strong enrichment of immune cells (CD45; magenta) and tumor cells (PanCK, green). Concordance between the proteogenomic assay and single analyte (**B**) protein and (**C**) RNA controls. Concordance between the initial and stored proteogenomic slide for (**D**) protein and (**E**) RNA analytes. (**F**) Multiplexed protein and RNA characterization of CRC sample with representative images highlighting the segmentation of 300 μm circular ROIs into tumor (PanCK^+^) and immune (CD45^+^) enriched regions. Segments illuminated in white were collected, black regions were not. Protein and RNA counts were SNR transformed and protein targets with SNR > 3 and WTA RNA targets with SNR > 4 were used in the analysis. Unsupervised hierarchical clustering of detected (**G**) RNA and (**H**) protein targets for CRC. (**I**) Combined volcano plot of Protein and RNA expression in CRC. A subset of differentially expressed genes are labeled with colors matching their analyte.

We were also interested in the ability to profile CRC samples that had been stored in 1X TBS-T at 4°C, protected from light for one-week post-initial DSP collection. One of the key benefits of using the GeoMx DSP platform for spatial profiling is that it entails a non-destructive process where tissue sections can be stored and reprofiled on additional ROIs (23). In this experiment, each sample slide contained two FFPE CRC tissue serial sections, one of which was used in the initial DSP profiling. We matched ROIs from the initial collection onto the unprofiled section without a stripping/reprobing process. As expected, profiling of the SPG assay slide post-one-week storage provided a high correlation to the initial profiling data for both the protein (*R* = 0.97) and RNA (*R* = 0.80) analytes (Fig. 3D and E). Together, we demonstrated high concordance between ROIs across serial sections and slides; more importantly, samples can be stored for additional DSP profiling runs.

### Optical Segmentation with Spatial Proteogenomic Assay

We then explored the optical dissection capabilities of the GeoMx segmentation function using the SPG assay. The ability to optically dissect a tissue based on morphological or biological features is an advantage for profiling protein and RNA targets in the distinct spatial context and within specific cell subpopulations. Using the segmentation strategies of the GeoMx DSP, we assessed the performance of the SPG assay on human CRC tissue samples. Applying the established SPG assay workflow, tissue sections were stained with the GeoMx Human WTA probe set and the 147-plex modular GeoMx Human NGS Protein panel, along with fluorescent-dye-conjugated primary antibodies for CD45 and PanCK. Twelve circular ROIs of 300 μm in diameter were segmented into the AOIs of CD45-enriched immune and PanCK-enriched tumor cell subpopulations. The ROIs were selected across the various tumor regions within the tissue section, including tumor regions proximal to immune-rich or immune-poor regions (Fig. 3F and Supplementary Fig. S9B). To analyze expression patterns within the segmented AOIs, we performed unsupervised hierarchical clustering of detected RNA and protein targets. As expected, we observed distinct clustering of targets within immune segments and tumor segments (Fig. 3G and H).

We performed differential expression analysis between CD45-enriched immune and PanCK-enriched tumor segments for both analytes. As shown in the volcano plot (Fig. 3I) we observed robust co-detection of immune-related protein and RNA targets in the CD45 enriched segments whereas tumor associated targets were observed in PanCK-enriched segments. In addition, an examination of expression levels of key RNA/Protein target pairs associated with either immune or tumor segments illustrates the variability of certain targets in a distinct immune or tumor AOI (Supplementary Fig. S11). For example, we observed high levels of HLA-DR protein/RNA target pair in immune segments and low in tumor. The opposite was true for cytokeratins, where high levels were observed in tumor segments and low levels in immune. Some RNA/protein target pairs, for example RPS6, a critical component to translational activities, were expressed at high levels in all segments.

When considering the correlations of all detectable protein targets and RNA targets in either immune or tumor segmented AOIs, we noted distinct patterns of positive and negative correlation. For example, interferon γ-inducible protein 30 (IFI30) is expressed in most antigen-presenting cells (APCs), including monocytes, macrophages, and dendritic cells, where it functions in MHC class II-restricted antigen processing and has been shown to have a role in the immune response to malignant tumors such as melanoma, prostate cancer, and glioma (33–36). In the immune AOIs, there was a strong positive correlation between the *IFI30* gene and protein targets associated with MHC class-II presentation on macrophages (CD68, CD14, HLA-DR), helper T-cells (CD4, CD127), dendritic cells (S100B) and tumor cells or activated APCs (B7-H3) (Supplementary Fig. S12). On the contrary, we observed a negative correlation of the *IFI30* gene with CD20 (B cells) and CD95/Fas (cell death) and CD8 (cytotoxic T cells) (Supplementary Fig. S12). In the tumor segments, negative correlation between *MUC5AC* RNA target associated with mucus production in goblet cells, and the protein target PanCK (tumor cell marker) was observed (37) (Supplementary Fig. S13). Conversely, there was a positive correlation between the *MUC5AC* RNA target and autophagy-related protein targets ATG5, ATG12, LAMP2A, and BAG3. Normal regulation of mucus production commonly involves autophagy for the regulation and secretion of mucins (38,39). Abnormal expression of MUC5AC is commonly associated with malignant colorectal cancerous cells (37), as observed in our data (Supplementary Fig. S13).

In addition to CRC tissues, we also evaluated the GeoMx segmentation capabilities on human NSCLC samples. Tissue sections were stained with the GeoMx Human WTA and the 147-plex modular GeoMx Human NGS Protein panel, along with primary antibodies for CD45 and PanCK for visualization. Twelve circular ROIs of 300 μm in diameter were selected across various tumor regions and segmented into CD45-enriched immune and PanCK-enriched tumor subpopulations (Supplementary Fig. S14A). As with CRC, the differential expression analysis demonstrated robust co-detection and specificity of both protein and RNA targets within each segment type (Supplementary Fig. S14B). Furthermore, we observed distinct clustering of RNA and protein targets within immune segments and tumor segments (Supplementary Fig. S14C).

### Performance of Spatial Proteogenomic Assay on Mouse Tissue

Given the frequent use of mouse models for studying various human biology and diseases, we examined our SPG assay on mouse tissue samples. For profiling, FFPE mouse multi-organ tissue microarray (TMA) sections were stained with the GeoMx Mouse WTA (21,000+ protein-coding genes) and the 15-modular GeoMx Mouse NGS Protein panel (137-plex) (Supplementary Table S4). In parallel, TMAs were stained with the standard single-analyte 137-plex GeoMx Mouse NGS Protein panel or with the standard single-analyte GeoMx Mouse WTA, used as the protein and RNA controls, respectively. For these experiments, circular ROIs (100 μm diameter) were selected for multiple tissue types including thymus, small intestine, kidney, and seminal vesicle (Fig. 4A, middle panels, and Supplementary Fig. S15). In each tissue type, there was a fairly strong correlation between the single-analyte RNA data and the SPG assay data on the small intestine (*R* = 0.84), thymus (*R* = 0.79), kidney (*R* = 0.81), and seminal vesicle (*R* = 0.83) (Fig. 4A, top panels). When we performed unsupervised hierarchical clustering on matched AOIs between the SPG assay and single-analyte RNA data, there was a high concordance between matched ROIs as well as a high correlation between tissue-specific ROIs (Supplementary Fig. S16A). In the unsupervised hierarchical clustering analysis of the top 500 detected RNA targets, we observed distinct clustering with respect to tissue type (Fig. 4B).

**Figure 4:**
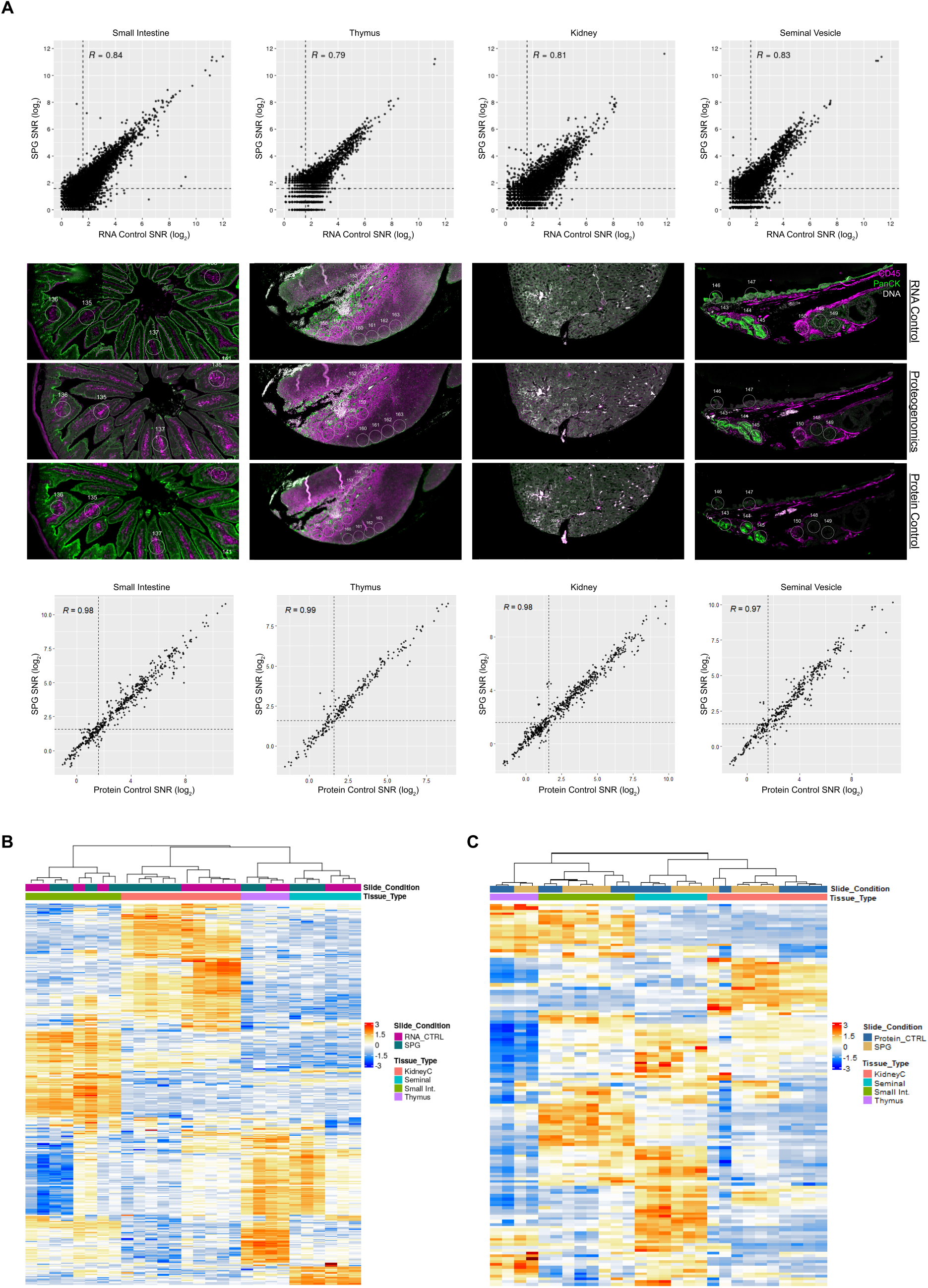
Spatial Proteogenomics across Mouse Tissue types. High plex spatial proteogenomic characterization of Mouse tissues with circular matched ROIs. FFPE sections were stained with GeoMx Mm WTA (RNA control), 15 stacked GeoMx Mouse Protein Modules (137-plex) (protein control), or both analytes simultaneously with the spatial proteogenomic workflow. (**A**) Concordance and representative images of mouse tissue used in the assessment of the spatial proteogenomic versus the respective (top) RNA and (bottom) protein control. Unsupervised hierarchical clustering of (**B**) top 500 RNA targets and (**C**) protein targets across all tissue types.

For protein profiling, the correlation between the SPG assay data to the single-analyte protein data was strong (*R* > 0.97) for all mouse tissue types tested (Fig. 4A, bottom panels). Hierarchical clustering analysis of the SPG assay and the single-analyte protein data was also performed on all detected protein targets and on matched ROIs in each tissue type. In an unsupervised hierarchical clustering on matched ROIs between the SPG assay and single-analyte protein data, there was a high concordance between matched ROIs as well as a high correlation between tissue-specific ROIs (Supplementary Fig. S16B). Furthermore, we observed distinct clustering of protein targets within specific tissue types (Fig. 4C). When we examined the expression level of several RNA/protein target pairs, there was a high concordance between the two analytes for GAPDH, S6, Histone H3, whereas a subset of targets was detectable only at the protein level (B7-H3, CD34, CD4, CD44, CD68, GZMB, S100B, SMA) (Supplementary Fig. S17).

### Glioblastoma Study using Spatial Proteogenomic Assay

After successfully developing a spatial proteogenomic assay and evaluating its performance on various cell lines and tissues, we assessed the performance of the SPG assay in another biological context using a brain glioblastoma tissue array. Glioblastoma multiforme (GBM) is a highly aggressive, grade IV astrocytoma that accounts for 49% of all primary malignant brain tumors (40,41). Despite low incidence compared to other human cancers, glioblastomas are considered one of the deadliest (41,42). The low numbers of patients coupled with high intra- and inter-tumor heterogeneity have presented an obstacle to developing glioblastoma treatments or providing formal subtypes that could contribute to therapeutic understanding (43). Initially, even cytological or immunohistochemical characteristics were difficult to solidify (44). The advancement of technology, however, has expanded knowledge of the molecular and genetic hallmarks of glioblastoma subtypes. The cytological hallmark of any GBM subtypes is the result of several mutations acting in concert. Additionally, overactivity of both PI3K/AKT and MAPK/ERK pathways drives cell proliferation and differentiation, a characteristic of GBM (45). It is also clear that these features have a variety of effects on prognosis (46,47). Giant cell glioblastomas (gcGBM) are of particular interest, as they are consistently associated with complete resection and improved survival (48). As another hallmark of gcGBM, giant, multinucleated cells have been linked to distinct, dysfunctional aspects of DNA damage signaling (49) and cell cycle checkpoints (50). Though many promising molecular targets for therapy have been identified (37–41), a better understanding of the tumor microenvironment and the tumor heterogeneity is needed for the development of more effective, and targeted therapies (51–55).

A spatial proteogenomic approach offers a robust way to identify impactful interactions between analytes. In this study, we profiled 23 different cases across 4 different pathological (grade 4) subtypes including glioblastoma multiforme (GBM; 17 cases), epithelioid glioblastoma (Ep-GBM; 3 cases), giant cell glioblastoma multiforme (gcGBM; 2 cases) and primitive neuronal components of glioblastoma (GBM-PNC; 1 case). The tissue microarray was stained with the GeoMx Human WTA and 147-plex modular GeoMx Human NGS Protein panel along with fluorophore-conjugated primary antibodies for CD45, and GFAP to guide ROI selection and segmentation. A total of 76 circular ROIs, 300 μm in diameter, were segmented into CD45^+^ (Immune), GFAP^+^ (Astrocyte/Tumor), or CD45^-^/GFAP^-^ (Fig. 5A and Supplementary Fig. S18). GBM is a highly heterogeneous and disorganized tumor, where more advanced tumor types are more disorganized. The heterogeneous and disorganized nature of GBM makes it difficult to select clear tumor- and/or immune-enriched ROIs compared to CRC or NSCLC samples. Even in these highly disorganized samples, the segmentation capabilities of the GeoMx DSP allowed us to optically dissect immune and astrocytic/tumor cells (Fig. 5A, 7A and Supplementary Figs. S18, S19). In total, 224 AOIs were collected; the resulting dataset was composed of 4.25 million data points (Fig. 5B).

**Figure 5:**
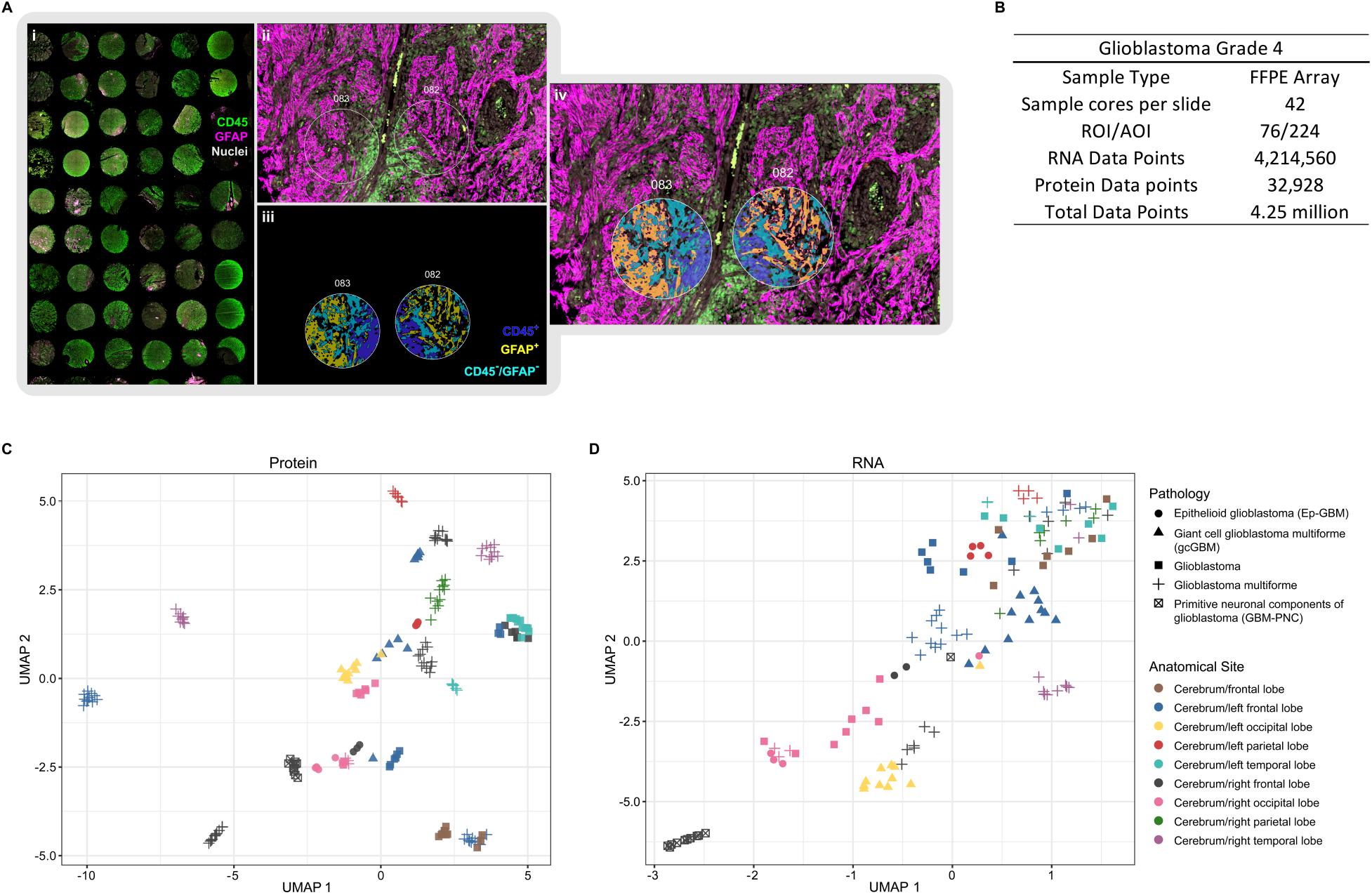
Spatial Proteogenomic exploration of Glioblastoma multiforme Grade 4. **(A)** 42 cores across 23 distinct sample sources and multiple GBM types. ROI were segmented into CD45^+^, GFAP^+^, or CD45^-^/GFAP^-^. (**B**) Statistics from a single slide and single GeoMx Spatial Proteogenomic run. UMAP (Uniform Manifold Approximation and Projection) plots for (**C**) protein and (**D**) RNA analytes.

In a uniform manifold approximation and projection (UMAP) dimension reduction analysis of GBM samples, we observed distinct clustering based on pathology and tumor anatomic location for both protein (Fig. 5C) and RNA (Fig. 5D). These findings suggest that the anatomic location and subtype of GBM contribute to overall pathology and the presence of molecular signatures. In a differential expression analysis between GFAP^+^ (Astrocyte/Tumor) and CD45^+^ (immune) segments using a cutoff of FDR < 0.05, and fold change (FC) > 2, we identified 25 proteins and 67 genes that were differentially expressed in a subset of samples.

The volcano plot highlighted the differential expression of key targets for both protein and RNA (Fig. 6A). For example, in CD45-enriched segments, we noted the higher expression of key protein targets associated with immune response such as T-cell (CD4), macrophage/monocytes (CD68, CD163, CD14, CD11c), and dendritic cells (CD11c, HLA-DR). In GFAP-enriched segments, tumor-associated proteins, such as CD56, CD44, and Tau (including phosphorylated variants S199, T231, and S404) were highly expressed (Fig. 6A) (56–58). Similarly, we observed higher levels of mRNA transcripts associated with immune response, such as macrophages *(SRGN, C1QA, C1QB, C1QC, IFI30)* and neutrophils *(LCP1, LAPTM5),* in CD45-enriched segments, whereas GBM-associated genes such as *GFAP, DDR1, CRYAB, SOX2, TTYH1* were expressed higher in GFAP-enriched segments (Fig. 6A) (58–65). Unsupervised hierarchical clustering analysis revealed distinct clustering of RNA (Fig. 6B) and protein (Fig. 6C) targets with respect to pathology as well as anatomic location. These observations are consistent with the UMAP analysis, showing that the anatomic location of tumors contributes to the molecular signatures of GBM subtypes.

**Figure 6:**
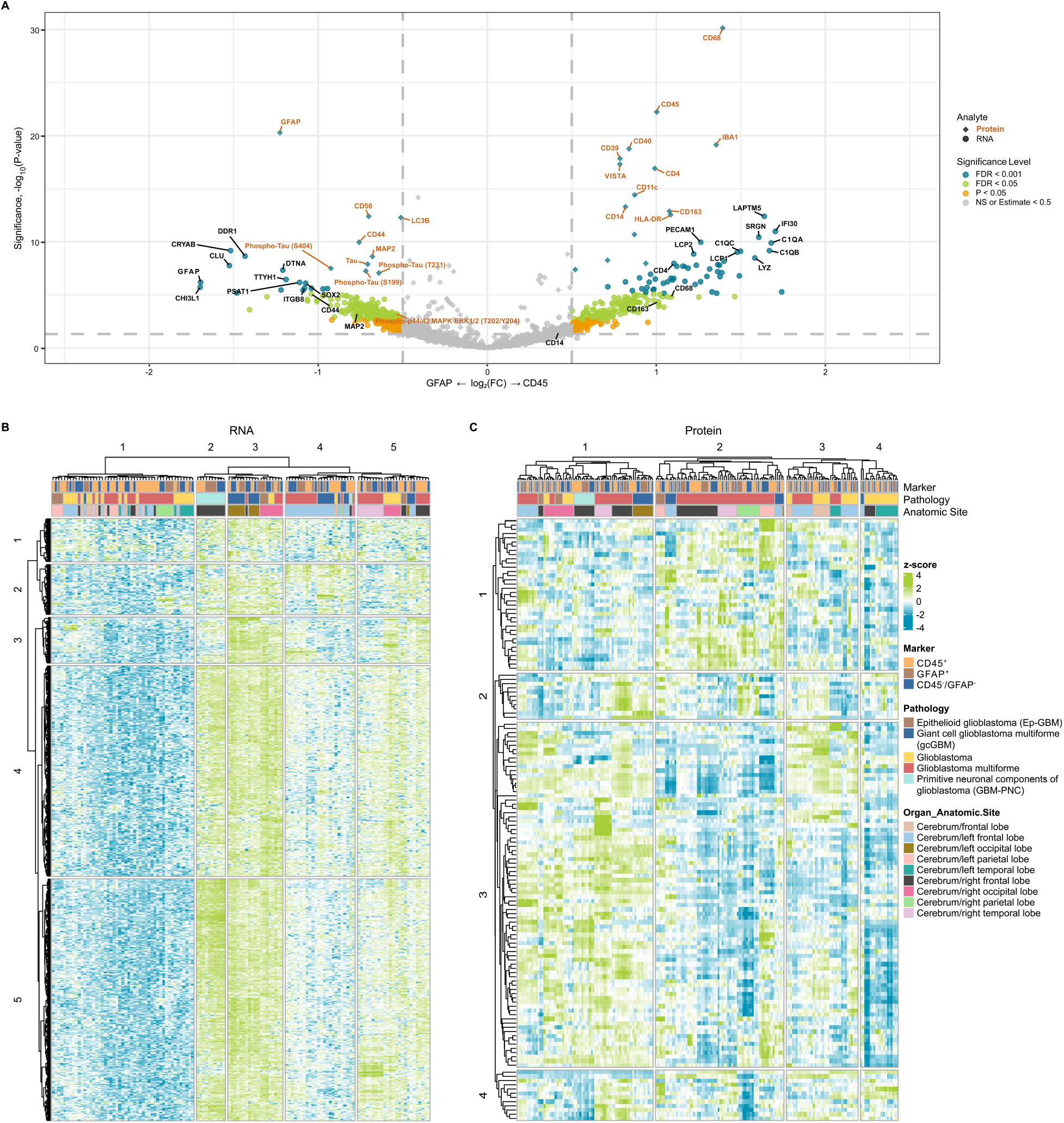
Differential expression analysis between GFAP and CD45 enriched segments. (**A**) Combined volcano plot resulting from the differential expression analysis between GFAP and CD45 enriched segments for RNA (•) and protein (♦) analytes. Targets with significantly differential expression are highlighted either orange (P-value < 0.05), green (FDR < 0.05) or blue (FDR < 0.01); whereas targets in grey show no significant difference in expression. A subset of differentially expressed targets are labeled. Unsupervised hierarchical clustering analysis of detected (**B**) RNA and (**C**) protein.

As noted above, GBM can be classified into several subtypes, one of which is gcGBM. In this study, one of our interests was identifying transcriptomic and proteomic signatures that differentiate gcGBM from GBM. To mitigate the influence of the tumor anatomical location on these different pathologies, we focused our analysis on left frontal lobe tumors, as a result, the analysis was based on a limited number of patient cases, one gcGBM and two GBM samples. Each case contained two tissue cores and two ROIs per core which were segmented into 3 AOIs: CD45^+^, GFAP^+^, or CD457^-^/GFAP^-^. In a differential expression analysis (FDR < 0.05, FC > 2) on detected targets for each AOI within the left frontal lobe, we identified a total of 64 and 5 differentially expressed genes and proteins, respectively between the two pathologies (Supplementary Fig. S20). For protein data, we also performed a differential expression analysis between GBM and gcGBM for CD45 (Supplementary Fig. S21A) or GFAP (Supplementary Fig. S21B) enriched segments. Using a cutoff of *P* < 0.05, FDR < 0.01, and FC ≥ 2, we identified a total of 14 and 7 differentially expressed proteins in CD45-and GFAP-enriched segments, respectively.

While the limited number of patient cases used in our analysis is insufficient for the complete elucidation of subtype-specific molecular signatures, we did observe several genes and proteins in gcGBM with distinct expression profiles compared to GBM (Fig. 7, Supplementary Fig. S20 and S21). For example, we observed differential expression of protein targets, CD3 and CD8, associated with infiltrating total T and cytotoxic T lymphocytes, respectively (Fig. 7B, Supplementary Fig. S21A). Immune infiltration has been noted as a characteristic of gcGBM (53,66,67). As the prognostic role of CD3 and CD8 have yet to be established in gcGBM, there are numerous reports associating infiltrating lymphocytes with improved prognosis for several cancers (68–71). In our analysis, both CD3 and CD8 proteins were expressed at least two-fold higher in several CD45-enriched AOIs of gcGBM compared to the same type of AOI in GBM samples (Fig. 7B). At the transcript level, *CD3E* and *CD8A* did not appear to be significantly differentially expressed between the two pathologies (Fig. 7C, Supplementary Fig. S20).

**Figure 7:**
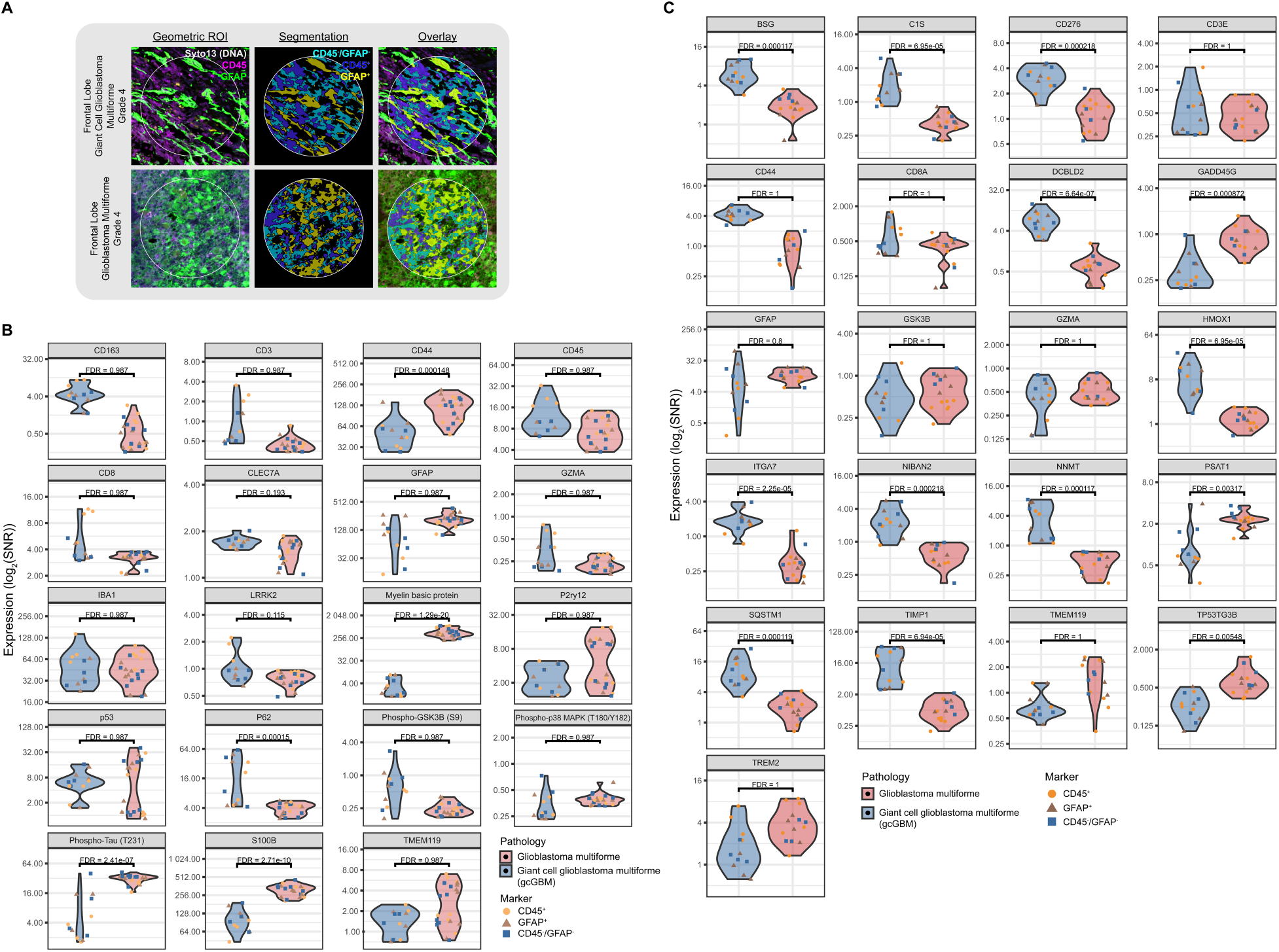
Spatial Proteogenomic exploration of gcGBM and GBM. (**A**) Representative images of ROIs segmented into CD45^+^, GFAP^+^, or CD457^-^/GFAP^-^. Top differentially expressed (**B**) Protein Targets and (**C**) RNA Targets from Frontal Lobe Giant Cell Glioblastoma Multiforme as compared to Glioblastoma Multiforme, all Grade 4.

We also evaluated several targets associated with tumor proliferation and/or migration to discriminate gcGBM from GBM. *GFAP,* for example, shows differential RNA expression with a lower average (and wider range) of SNR in gcGBM than GBM samples (Fig. 7C). This difference in expression between the two pathologies is maintained in GFAP protein (Fig. 7B), which is known to positively correlate with advancing neuroglial tumor grade (72).

Enrichment of S100B has been shown to be sufficient to drive angiogenesis, increase tumor inflammation (73), enhance growth (74) and drive migration (75), all of which are features of GBM. The proposed tumorigenic contribution of S100B has been associated with its ability to attenuate the tumor suppressive activity of p53 by inhibiting p53 phosphorylation and regulating cell proliferation and differentiation through increasing mitogenic kinase Ndr (76) and Akt (77) activity (78). *In vitro* studies involving cell lines derived from malignant glioblastoma shown that the attenuation of S100B expression decreased the number of invading cells whereas upregulation resulted in an increase in invading cells (79). In our analysis, S100B protein expression in gcGBM was decreased at least 2-fold in all AOI as compared to GBM (Fig. 7B, Supplementary Fig. S21A and S21B). While it has been reported that higher expression levels of S100B in GBM patients is associated with shorter overall survival (79), it is unclear what role S100B may play in the favorable prognosis of gcGBM compared to GBM.

While CD44, at the protein level, exhibited a broad range of expression for both pathologies, the average expression level of RNA was at least 2-fold higher in gcGBM than that of GBM (Fig. 7B and 7C). CD44 expression is critical for GBM invasion and migration; more importantly, GBM cells with higher levels of CD44 have been associated with poor prognosis (56,57,80,81).

In our analysis, the average expression level for MBP protein in gcGBM was at least 64 times lower in all AOIs compared to GBM samples (Fig. 7B, Supplementary Fig. S21A and S21B). On the other hand, differential expression at the transcript level was not observed. Myelin basic protein (MBP) has been viewed as a potential biomarker for various types of tumors including glioblastoma in cerebrospinal fluids (82–84). In other studies, MBP was highly expressed in oligodendroglioma while minimally expressed in GBM (85). It has been suggested that MBP may play a role in the progression and potential treatment of multiple sclerosis (MS), however, it is unclear what role it has in gcGBM biology (86).

As noted earlier, the phosphorylation state of proteins can only be captured through protein and not transcriptomic analysis. GSK3β is part of the PI3K-AKT/mTOR pathway whose phosphorylation state plays an important role in glycogen synthesis, apoptosis, angiogenesis, and cell cycle. In our dataset, protein expression levels of phospho-GSK3β (Ser9) were broad and at least 2-fold higher for several AOIs in the gcGBM sample compared to GBM sample AOIs (Fig. 7B). Phosphorylation at Ser9 of GSK3β leads to its inactivation, which in turn prevents the phosphorylation and subsequent degradation of β-catenin. Active β-catenin has been shown to drive cell proliferation in GBM (87–89). We also observed differential protein expression phosphorylated Tau variants. Phospho-Thr231 Tau was at least 2-fold higher for several AOIs in the GBM sample compared to gcGBM (Fig. 7B, Supplementary Fig. S21A and S21B). Whereas Tau, pSer199 Tau and pSer404 Tau were only expressed at higher levels in GBM than gcGBM within the GFAP enriched AOIs (Supplementary Fig. S21B). While associated with neurodegenerative diseases such as Alzheimer’s disease, altered Tau expression has been observed in several cancers including glioblastoma (90–95). Like with the other targets mentioned above, it is not clear based on our limited data set what role of these phosphorylated proteins play in GBM cell migration, survival, and apoptosis. Though limited in size, our study exemplifies the utility of the SPG assay and its potential contribution in expanding our understanding of GBM molecular pathology.

## Discussion

To better understand the relationship between RNA and protein within a spatially defined cell population, we have developed a high-plex, GeoMx DSP Spatial Proteogenomic assay with NGS readout to simultaneously detect RNA (whole transcriptome, > 18,000-plex) and protein (> 100-plex) from a single tissue section slide. We have confirmed that the sensitivity and specificity for both analytes under SPG assay conditions are comparable to single analyte conditions where only a slight loss of sensitivity (< 11%) is observed for both analytes. The slight loss in sensitivity is a tradeoff when implanting the SPG assay given that the two analytes required two distinct and disparate antigen retrieval conditions for optimal detection. We have highlighted several use cases in cell lines and tissue derived from human and mouse to demonstrate how this workflow can be leveraged to accurately measure RNA and protein within the same sample. Additionally, we demonstrate simultaneous high-plex detection of distinct immune or tumor RNA and protein targets from spatially resolved individual cell subpopulations in human CRC and NSCLC using the tissue segmentation capabilities of the GeoMx DSP with the SPG assay. Unlike CRC and NSCLC, GBM is a highly heterogeneous and disorganized tumor, where selecting the clearly defined immune- and/or tumor-enrich regions can be challenging. However, using the GeoMx DSP we were able to optically dissect immune and astrocytic/tumor cells in various GBM subtypes. While the GBM study was restricted in scope due to the limited number of case samples, we did observe distinct clustering of both RNA and protein based on pathology and anatomic location. Furthermore, we were able to identify the differential expression between GBM and gcGBM for several protein and RNA targets.

Overall, the SPG assay expands upon the capabilities of the GeoMx DSP and enables researchers to conduct a comprehensive molecular analysis while preserving precious samples. Profiling RNA or protein alone will only provide a limited picture of the biological system under study, especially when the correlation between RNA expression and protein abundance can be poor, as highlighted in several of our case use studies. We also identified several targets that could only be detected by protein and not by RNA. One of the key advantages of implementing the SPG assay is that it enables the profiling of both RNA and protein from identical cell populations while eliminating the technical variation *(i.e.,* section-to-section variation and precisely matching ROIs across multiple samples) introduced when performing single-analyte assays. More importantly, the SPG assay provides an efficient means of profiling two analytes from a single slide as opposed to running two separate slides, one for each analyte. Another advantage of the SPG workflow, as with all GeoMx assays, is that the process is non-destructive which allows the researcher to store and reprofile their samples. Furthermore, it uses modular GeoMx reagent panels that are validated for use in high-plex studies and can be customized with additional targets. More importantly, GeoMx Protein reagents contain antibodies specific to intracellular and phosphorylated protein targets. The detection and quantification of phosphorylated proteins can only be achieved at the protein and not RNA level by using phospho-specific antibodies. We have shown that the GeoMx Spatial Proteogenomic assay in FFPE tissues across species gives comparable specificity to the single-analyte GeoMx assays, with only a minor decrease in overall sensitivities, demonstrating a high-quality multi-modal omic workflow compatible with the GeoMx DSP platform.

Transcriptomics and proteomics work in concert to describe cell activity, cell function, and cell-to-cell communication. Profiling RNA expression gives insight into the cellular blueprints and assessing proteins describes the cellular architecture, functions, and cell-to-cell communications. But each in isolation is only part of the necessary whole. The ability to profile both RNA and protein analytes, at ultra high-plex within a spatially resolved single population of cell types, help to close the knowledge gap between these two omics, paving the way for critical spatial proteogenomic investigations and discoveries within the rapidly emerging field of spatial biology.

## Materials and Methods

All experiments were carried out using reagents recommended or provided by NanoString Technologies and are summarized in Supplementary Tables S1 and S2, respectively. A list of antibodies making up each of the Human and Mouse GeoMx Protein Modules are summarized in Supplementary Tables S3 and S4, respectively. All reagents and instruments are for Research Use Only (RUO) and not for use in diagnostic procedures.

### FFPE Samples

Sections of formalin-fixed, paraffin-embedded (FFPE) cell pellet arrays (CPAs) and tissues, 5 μm in thickness, were used in these studies. Two custom CPAs consisting of 11 and 45 human cell lines were generated by Acepix (Hayward, CA); cell lines making up the CPA can be found in Supplementary Tables S5 and S6. FFPE sections of human colorectal cancer (CRC) were from either BioChain (cat #: T22235090-1) or ProteoGenex (cat#: 025562T2). Human FFPE non-small cell lung cancer (NSCLC) was from US BioMax (cat #: HuCAT231) or ProteoGenex (cat#: 041556T2). Mouse multi-organ tissue microarray (TMA) was from SuperBiochips (cat#: ZE1) and human brain glioblastoma tissue array was from US Biomax (cat#: GL806g).

### GeoMx Single Analyte Protein and RNA Assay

For protein only control, slides were manually processed according to the Protein FFPE Manual Slide Preparation Protocol in the GeoMx-NGS Slide Preparation User Manual (MAN-10150) and associated published material (23).

RNA only control slides were processed according to the RNA FFPE BOND RX Slide Preparation Protocol in the GeoMx-NGS Slide Preparation User Manual for FFPE (MAN-10151) and associated published materials (22,23,25).

### GeoMx Spatial Proteogenomic Assay Sample Prep

Spatial proteogenomic slides were processed according to the GeoMx DSP Spatial Proteogenomic Protocol (MAN-10158).

### GeoMx DSP Experiments ROI Selection and Collection

For the spatial proteogenomic slides, GeoMx Digital Spatial Profiling was carried out according to GeoMx-DSP Spatial Proteogenomic Protocol (MAN-10158); whereas the control slides were processed according to GeoMx-NGS DSP Instrument User Manual (MAN-10152) and as described by Merritt, et al. (23,25). For CPAs, two geometric regions of interest (ROIs) of 200 μm in diameter were profiled per cell line.

Tissue sections were stained with fluorescent morphology markers and nuclear counterstain (Syto-13) to aid in the selection of areas of interest (AOIs). Human CRC, human NSCLC, and a mouse multi-organ tissue array were stained with the GeoMx Solid Tumor TME Morphology Kit (NanoString, GMX-PRO-MORPH-HST-12 and GMX-RNA-MORPH-HST-FFPE-12, for human and mouse, respectively) to aid in the visualization of immune (CD45^+^) and epithelial/tumor (PanCK^+^) enriched regions. Human brain glioblastoma with normal brain tissue microarray was stained with the GeoMx Solid Tumor TME Morphology Kit and anti-GFAP (astrocytes).

In human tissues, circular geometric ROIs of 100 μm in diameter were collected for each morphology marker-specific ROI. In the mouse multi-organ array, anatomically distinct regions were selected for specific tissue and then sampled with 100 μm diameter circular geometric ROIs. Two to three regions were selected for each tissue. For each tissue type, ROIs were matched across all test slides under study. An advanced ROI selection strategy (segmentation) was implemented on CRC, NSCLC, and GBM. For segmentation experiments, circular ROIs of 300 μm in diameter were segmented into marker-specific areas of interest using the GeoMx auto-segmentation tool (26).

### Next-Generation Sequencing and Data Analysis

Library preparations were carried out according to the GeoMx-DSP Spatial Proteogenomic Protocol (MAN-10158). Libraries were sequenced on an Illumina NextSeq2000 or NovaSeq6000 according to the manufacturer’s instructions.

The resulting FASTQ files were processed along with a modified GeoMx NGS Pipeline config file using the NanoString GeoMx NGS Pipeline v2.0 or v2.3 according to the GeoMx DSP NGS Readout User Manual (MAN-10153). As recommended in MAN-10158, proteogenomic data was processed and analyzed using GeomxTools (v3.1.1; https://github.com/Nanostring-Biostats/GeomxTools/) (27) in R separately for the protein and RNA analytes. AOIs with low reads sequenced, low saturation, and low area were removed from analysis.

For RNA, the signal-to-noise ratio (SNR) was calculated by dividing the signal by the limit of quantitation (LoQ), where the LoQ is calculated as follows:

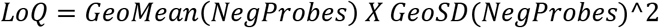

Gene counts were normalized by quartile 3 (Q3) normalization after the removal of genes below the detection threshold. For tissue, genes were filtered to those above the detection threshold in >15% of AOIs. For protein, the SNR was calculated by dividing the signal by the geomean of the three IgG negative controls (mouse IgG1 and IgG2a, and rabbit IgG).

Cluster heatmaps were generated with the pheatmap R package (28). Clustering was carried out on log transformed scaled counts using the pheatmap “correlation” method. For tissue, differential expression analysis between different populations of cells was performed for each analyte using a two-sided, unpaired *t*-test from rstatix R package. The threshold for significance was set at *p-* value < 0.05 and adjusted for multiple comparisons (or multiple hypothesis testing) using the Benjamini-Hochberg method (29). The estimated fold changes (log2FC) in the SNR and *p*-values for both analytes were summarized in a single volcano plot using R ggplot2 package.

### Advanced Data Analysis

Proteogenomic data generated from the GBM studies were processed and analyzed using GeomxTools (v3.1.1) in R separately for the protein and RNA analytes as noted above. For the protein analyte, proteogenomic data underwent background normalization using the geometric mean of three IgG negative controls. For RNA, we calculated the LOQ for each AOI and removed segments where less than 5% of the genes were expressed above LOQ. Using LOQ, we then removed genes that were not detected in at least 10% of the ROIs. This final, filtered dataset underwent Q3 normalization and used for further analysis.

### Statistical analyses

All analyses were performed in R (v 4.1.2). UMAP (Uniform Manifold Approximation and Projection) plots were generated with normalized expression data using the umap packages (v0.2.8.0) using default settings. Coefficients of variation (CV) were calculated for each protein or gene (*CV_g_*= *SD_g_/mean_g_*). The genes with the highest CVs were filtered and plotted as a heatmap using unsupervised hierarchical clustering based on Pearson distance. Heatmaps were generated using the ComplexHeatmap (30) package (v2.10.0).

Differential expression (DE) analysis was performed on a per-gene basis where the normalized expression was modeled using a linear mixed-effect model (LMM) to account for multiple sampling of ROI/AOI segments per tissue. DE was performed using the mixedModelDE function from the GeomxTools package (v3.1.1). DE results were visualized in volcano plots and violin plots using the ggplot2 package (v 3.3.6). Volcano plots are used to visualize the overall results of DE with estimated fold changes (log2FC) in the SNR and *p*-values plotted for each contrast. Violin plots highlight features of interest and the most differentially expressed genes or proteins in a comparison, visualized using SNR. For protein data, SNR is equal to the signal of a protein divided by the geometric mean of the three background probes. For RNA expression data, SNR is equal to the signal divided by LOQ.

## Data Availability

All raw and processed data used in this manuscript will be available upon request in writing to the corresponding author.

## Acknowledgements

We thank Bridget Kulasekara, Karen Nguyen, Daniel R. Zollinger, Michelle Kriner, Michael Rhodes, Erin Piazza, and Margaret Hoang from NanoString Technologies for their insight and guidance. We also want to thank Corey Williams from NanoString for their help with NGS.

