## Supplementary Table 1 for "Ultra High-Plex Spatial Proteogenomic Investigation of Giant Cell Glioblastoma Multiforme Immune Infiltrates Reveals Distinct Protein and RNA Expression Profiles"

| Reagent/Material | Source/Part Number |
|---|---|
| 10% neutral buffered formalin (NBF) | EMS Diasum, 15740-04 |
| 100% deionized formamide | VWR, VWRV0606 |
| 100% ethanol (EtOH): ACS grade or better | Various |
| 10X citrate buffer pH 6 | Sigma-Aldrich, C9999-1000ML |
| 10X phosphate buffered saline pH 7.4 (PBS) | Invitrogen, 00652158 |
| 10X TBS with tween 20 (TBS-T) | Cell Signaling Technologies, 9997S |
| 10X tris buffered saline (TBS) | Cell Signaling Technologies, 12498S |
| 16% paraformaldehyde (PFA) | 16% stock, 28908 |
| 1X phosphate buffered saline pH 7.4 (PBS) | ThermoFisher,10010031 |
| 20X SSC (DNase, RNase free) | Sigma-Aldrich, S6639 |
| BOND Dewax Solution, 1 L | Leica Biosystems, AR9222 |
| BOND Epitope Retrieval 1, 1 L | Leica Biosystems, AR9961 |
| BOND Epitope Retrieval 2, 1 L | Leica Biosystems, AR9640 |
| BOND Wash Solution 10X Concentrate, 1 L | Leica Biosystems, AR9590 |
| CitriSolv | Fisher Scientific, 04-355-121 |
| DEPC-treated water | ThermoFisher, AM9922 |
| Glycine | Sigma-Aldrich, G7126 |
| Proteinase K | Ambion, AM2546 |
| Tris base | Sigma-Aldrich, 10708976001 |
| HybriSlip hybridization covers (22 mm x 40 mm x 0.25 mm) | Grace Bio-Labs, 714022 |
| RNase Away | Thermofisher, 7003PK |
| Dextran sulfate | Sigma, 67578 |
| BSA | VWR, 97061-420 |
| Deoxyribonucleic acid, single stranded from salmon testes (denatured by heating at 95$^{o}$C for 10 min prior to use) | Sigma, D7656 |
| Agencourt AMPure XP | Beckman Coulter, A63880 |
| Elution buffer (Tris-HCl 10 mM with 0.05% Tween-20, pH 8.0 ) * | Teknova, T1485 |
| Bioanalyzer DNA High Sensitivity Kit | Agilent, 5067-4626 |
| Quibit dsDNA HS Assay Kit | Q32854 |
| GFAP antibody (GA-5) Alexa Fluor(R) 647 | Novus Biologicals, NBP2-33184AF647 |
| nCounter Master Kit (Maxor FLEX Systems) Reagents and Cartridges | NAA-AKIT-012 |
| BOND Research Detection System (includes 6x 30 mL Open Containers) | Leica Biosystems, DS9455 |
| BOND Titration Kit (includes 50 inserts) | Leica Biosystems, OPT9049 |
| BOND Universal Covertiles | Leica Biosystems, S21.2001 |
| BOND Open Containers 30 mL | Leica Biosystems, OP309700 |

**Supplementary Table S1:** A list of materials and reagents that are recommended for the GeoMx DSP assays that are not supplied by NanoString.
