## Supplementary Table 2 for "Ultra High-Plex Spatial Proteogenomic Investigation of Giant Cell Glioblastoma Multiforme Immune Infiltrates Reveals Distinct Protein and RNA Expression Profiles"

| Reagent/Material | Source/Part Number |
|---|---|
| GeoMx Solid Tumor TME Morphology Kit Human Protein Compatible | NanoString, GMX-PRO-MORP-HHST-12 |
| GeoMx Immune Cell Profiling Panel Human Protein Core for nCounter | NanoString, GMX-PROCO-NCT-HICP-12 |
| GeoMx Immune Activation Status Panel Human Protein Module for nCounter | NanoString, GMX-PROMOD-NCT-HICT-12 |
| GeoMx Immune Cell Typing Panel Human Protein Module for nCounter | NanoString, GMX-PROMOD-NCT-HICT-12 |
| GeoMx IO Drug Target Panel Human Protein Module for nCounter | NanoString, GMX-PROMOD-NCT-HIODT-12 |
| GeoMx Protein Slide Prep Kit for FFPE | NanoString, GMX-PREP-PRO-FFPE-12 |
| GeoMx Hyb Code Pack Protein | NanoString, GMX-PRO-HYB-96 |
| GeoMx DSP Collection Plate | NanoString, GMX-DSP-COLL-PLT-4 |
| GeoMx DSP Instrument Buffer Kit | NanoString, GMX-DSP-BUFF-KIT |
| GeoMx Human Protein Core for NGS | NanoString, GMX-PROCO-NGS-HCORE-12 |
| GeoMx Immune Cell Typing Panel Human Protein Module for NGS | NanoString, GMX-PROMOD-NGS-HICT-12 |
| GeoMx IO Drug Target Panel Human Protein Module for NGS | NanoString, GMX-PROMOD-NGS-HIODT-12 |
| GeoMx Immune Activation Status Panel Human Protein Module for NGS | NanoString, GMX-PROMOD-NGS-HIAS-12 |
| GeoMx Pan-Tumor Panel Human Protein Module for NGS | NanoString, GMX-PROMOD-NGS-HPT-12 |
| GeoMx Myeloid Panel Human Protein Module for NGS | NanoString, GMX-PROMOD-NGS-HMY-12 |
| GeoMx MAPK Signaling Panel Human Protein Module for NGS | NanoString, GMX-PROMOD-NGS-HMAPK-12 |
| GeoMx PI3K/AKT Signaling Panel Human Protein Module for NGS | NanoString, GMX-PROMOD-NGS-HPI3K-12 |
| GeoMx Neural Cell Typing Panel Human Protein Module for NGS | NanoString, GMX-PROMOD-NGS-HNCT-12 |
| GeoMx Alzheimer's Pathology Panel Human Protein Module for NGS | NanoString, GMX-PROMOD-NGS-HADP-12 |
| GeoMx Seq Code Pack | NanoString, GMX-NGS-SEQ-AB |
| GeoMx Human Whole Transcriptome Atlas Human RNA for Illumina Systems | NanoString, GMX-RNA-NGS-HuWTA-4 |
| GeoMx RNA Slide Prep Kit for FFPE | NanoString, GMX-PREP-RNA-FFPE-12 |
| GeoMx Cancer Transcriptome Atlas Human RNA for Illumina Systems | NanoString, GMX-RNA-NGS-CTA-4 |

**Supplementary Table S2a.** A list of NanoString supplied materials and reagents.

| Reagent/Material | Source/Item Number |
|---|---|
| GeoMx Solid Tumor TME Morphology Kit Mouse RNA FFPE Compatible | NanoString, GMX-RNA-MORPH-MST-FFPE-12 |
| GeoMx Mouse Protein Core for NGS | NanoString, GMX-PROCO-NGS-MCORE-12 |
| GeoMx Immune Activation Status Panel Mouse Protein Module for NGS | NanoString, GMX-PROMOD-NGS-MIAS-12 |
| GeoMx IO Drug Target Panel Mouse Protein Module for NGS | NanoString, GMX-PROMOD-NGS-MIODT-12 |
| GeoMx Cell Death Panel Mouse Protein Module for NGS | NanoString, GMX-PROMOD-NGS-MCD-12 |
| GeoMx Immune Cell Typing Panel Mouse Protein Module for NGS | NanoString, GMX-PROMOD-NGS-MICT-12 |
| GeoMx Glial Cell Subtyping Panel Mouse Protein Module for NGS | NanoString, GMX-PROMOD-NGS-MGCS-12 |
| GeoMx Pan-Tumor Panel Mouse Protein Module for NGS | NanoString, GMX-PROMOD-NGS-MPT-12 |
| GeoMx Myeloid Panel Mouse Protein Module for NGS | NanoString, GMX-PROMOD-NGS-MMY-12 |
| GeoMx MAPK Signaling Panel Mouse Protein Module for NGS | NanoString, GMX-PROMOD-NGS-MMAPK-12 |
| GeoMx PI3K/AKT Signaling Panel Mouse Protein Module for NGS | NanoString, GMX-PROMOD-NGS-MPI3K-12 |
| GeoMx Neural Cell Typing Panel Mouse Protein Module for NGS | NanoString, GMX-PROMOD-NGS-MNCT-12 |
| GeoMx Alzheimer's Pathology Panel Mouse Protein Module for NGS | NanoString, GMX-PROMOD-NGS-MADP-12 |
| GeoMx Parkinson's Pathology Panel Mouse Protein Module for NGS | NanoString, GMX-PROMOD-NGS-MPDP-12 |
| GeoMx Alzheimer's Pathology Extended Panel Mouse Protein Module for NGS | NanoString, GMX-PROMOD-NGS-MADEP-12 |
| GeoMx Autophagy Panel Mouse Protein Module for NGS | NanoString, GMX-PROMOD-NGS-MA-12 |
| GeoMx Mouse Whole Transcriptome Atlas *Mouse RNA for Illumina Systems* | NanoString, GMX-RNA-NGS-MsWTA-4 |

**Supplementary Table S2b.** A list of NanoString supplied materials and reagents continued.
