## Supplementary Table 3 for "Ultra High-Plex Spatial Proteogenomic Investigation of Giant Cell Glioblastoma Multiforme Immune Infiltrates Reveals Distinct Protein and RNA Expression Profiles"

| NGS Human Panel | Target |
|---|---|
| GeoMx Human Immune Activation Status Panel | CD127 |
| | CD25 |
| | CD27 |
| | CD44 |
| | CD45RO |
| | CD80 |
| | ICOS |
| | PD-1 |
| | PD-L1 |
| | PD-L2 |
| GeoMx Human NGS Cell Death Panel | BAD |
| | BCL6 |
| | BCLXL |
| | BIM |
| | CD95/Fas |
| | Cleaved Caspase 9 |
| | GZMA |
| | NF-1 (Neurofibromin-1) |
| | p53 |
| | PARP |
| GeoMx Human NGS Immune Cell Typing Panel | CD20 |
| | CD3 |
| | CD34 |
| | CD4 |
| | CD56 |
| | CD66b |
| | CD8 |
| | Fibronectin |
| | FOXP3 |
| | GZMB |
| GeoMx Human NGS IO Drug Target Panel | 4-1BB |
| | B7-H3 |
| | CTLA4 |
| | GITR |
| | IDO1 |
| | LAG3 |
| | OX40L |
| | STING |
| | Tim-3 |
| | VISTA |
| GeoMx Human NGS MAPK Signaling Panel | BRAF |
| | EGFR |
| | JNK (phospho T183/Y185) |
| | MEK1 (phospho S217/S221) |
| | P38 (phospho T180/Y182) |
| | p44/42 MAPK ERK1/2 |
| | p44/42 MAPK ERK1/2 (phospho T202/Y204) |
| | P90RSK (phospho T359/S363) |
| | pan-RAS |
| GeoMx Human NGS Myeloid Panel | ARG1 |
| | CD11b |
| | CD11c |
| | CD14 |
| | CD163 |
| | CD39 |
| | CD40 |
| | CD68 |
| | HLA-DR |
| GeoMx Human NGS Pan-Tumor Panel | Bcl-2 |
| | EpCAM |
| | ER-alpha |
| | Her2 |
| | MART1 |
| | NY-ESO-1 |
| | PAN-CK |
| | PR |
| | PTEN |
| | SMA |
| GeoMx Human NGS PI3K/AKT Signaling Panel | AKT (phospho S473) |
| | GSK3 (phospho S9) |
| | GSK3αβ (phospho S21/S9) |
| | INPP4B |
| | MET |
| | Pan-AKT |
| | PLCG1 |
| | P-RAS-40 (phopho T246) |
| | TUBERIN (phospho T1462) |

| NGS Human Panel | Target |
|---|---|
| GeoMx Human NGS Alzheimer's Extended Pathology Panel | ADAM10 |
| | BACE1 |
| | IDE |
| | Neprilysin |
| | Neurogranin |
| | PSEN1 |
| | p-Tau (S199) |
| | p-Tau (S396) |
| | p-Tau (T214) |
| | p-Tau (T231) |
| GeoMx Human NGS Alzheimer's Pathology Panel | Amyloid Precursor Protein |
| | Amyloid-Beta 1-40 |
| | Amyloid-Beta 1-42 |
| | APOE |
| | P2RX7 |
| | Phospho-Tau (S404) |
| | Phospho-Tdp-43 (S409/S410) |
| | Tau |
| | Tdp-43 |
| | Ubiquitin |
| GeoMx Human NGS Autophagy Panel | ATG12 |
| | ATG5 |
| | BAG3 |
| | GBA |
| | HSC70 |
| | LAMP2A |
| | LC3B |
| | P62 |
| | TFEB |
| | VPS35 |
| GeoMx Human NGS Glial Cell Subtyping Panel | C4B |
| | CD9 |
| | Clec7a |
| | CSF1R |
| | Ctsd |
| | Emp1 |
| | GPNMB |
| | Mertk |
| | S100B |
| | VIM |
| GeoMx Human NGS Neural Cell Typing Panel | GFAP |
| | IBA1 |
| | MAP2 |
| | Myelin basic protein |
| | NeuN |
| | Neurofilament light |
| | Olig2 |
| | P2ry12 |
| | Synaptophysin |
| | TMEM119 |
| GeoMx Human NGS Parkinson's Pathology Panel | Alpha-synuclein |
| | ApoA-I |
| | Calbindin |
| | FUS |
| | LRRK2 |
| | Park5 |
| | Park7 |
| | Phospho-Alpha-synuclein (S129) |
| | PINK1 |
| | Tyrosine Hydroxylase |
| GeoMx Human NGS Core | Beta-2-microglobulin |
| | CD31 |
| | CD45 |
| | GAPDH |
| | Histone H3 |
| | Ki-67 |
| | Ms IgG1 |
| | Ms IgG2a |
| | Rb IgG |
| | S6 |

**Supplementary Table S3:** A list of Human protein targets by module.
