## Supplementary Table 4 for "Ultra High-Plex Spatial Proteogenomic Investigation of Giant Cell Glioblastoma Multiforme Immune Infiltrates Reveals Distinct Protein and RNA Expression Profiles"

| NGS Mouse Panel | Target |
|---|---|
| GeoMx Mouse Immune Activation Status Panel | CD127 |
| | CD27 |
| | CD40L |
| | CD44 |
| | CD86 |
| | ICOS |
| | PD-1 |
| | PD-L1 |
| GeoMx Mouse NGS Cell Death Panel | BAD |
| | BCLXL |
| | BIM |
| | Cleaved Caspase 3 |
| | gamma-H2AX |
| | NF-1 (Neurofibromin-1) |
| | p21 |
| | p53 |
| | PARP |
| | Perforin |
| GeoMx Mouse NGS Immune Cell Typing Panel | BatF3 |
| | CD19 |
| | CD28 |
| | CD3 |
| | CD34 |
| | CD4 |
| | CD8 |
| | Fibronectin |
| | FOXP3 |
| | GZMB |
| GeoMx Mouse NGS IO Drug Target Panel | B7-H3 |
| | CTLA4 |
| | GITR |
| | LAG3 |
| | OX40L |
| | Tim-3 |
| | VISTA |
| GeoMx Mouse NGS MAPK Signaling Panel | BRAF |
| | EGFR |
| | JNK (phospho T183/Y185) |
| | MEK1 |
| | MEK1 (phospho S217/S221) |
| | P38 |
| | p44/42 MAPK ERK1/2 |
| | p44/42 MAPK ERK1/2 (phospho T202/Y204) |
| | P90RSK (phospho T359/S363) |
| | pan-RAS |
| GeoMx Mouse NGS Myeloid Panel | CD11b |
| | CD11c |
| | CD14 |
| | CD163 |
| | CD39 |
| | CD40 |
| | CD68 |
| | F4/80 |
| | Ly6G/Ly6C |
| | MHC II |
| GeoMx Mouse NGS Pan-Tumor Panel | AhR |
| | AR |
| | EpCAM |
| | ER-alpha |
| | Her2 |
| | IFNGR |
| | PAN-CK |
| | Pmel17 |
| | SMA |
| GeoMx Mouse NGS PI3K/AKT Signaling Panel | AKT (phospho S473) |
| | AMPK-alpha_pThr172 |
| | GSK3αβ (phospho S21/S9) |
| | MET |
| | Pan-AKT |
| | PLCG1 |
| | P-RAS-40 (phopho T246) |
| | S6_pS235/236 |

| NGS Mouse Panel | Target |
|---|---|
| GeoMx Mouse NGS Alzheimer's Extended Pathology Panel | BACE1 |
| | IDE |
| | Neprilysin |
| | Neurogranin |
| | PSEN1 |
| | p-Tau (S199) |
| | p-Tau (S396) |
| | p-Tau (T214) |
| | p-Tau (T231) |
| GeoMx Mouse NGS Alzheimer's Pathology Panel | Amyloid Precursor Protein |
| | Amyloid-Beta 1-42 |
| | APOE |
| | P2RX7 |
| | Phospho-Tau (S404) |
| | Tau |
| | Tdp-43 |
| | Ubiquitin |
| GeoMx Mouse NGS Autophagy Panel | ATG12 |
| | ATG5 |
| | BAG3 |
| | Beclin-1 |
| | LC3B |
| | P62 |
| | PLA2G6 |
| | TFEB |
| | ULK1 |
| | VPS35 |
| GeoMx Mouse NGS Glial Cell Subtyping Panel | Aldh1l1 |
| | CD9 |
| | CSF1R |
| | Ctsd |
| | GPNMB |
| | Mertk |
| | MSR1 |
| | S100B |
| | SPP1 |
| | VIM |
| GeoMx Mouse NGS Neural Cell Typing Panel | GFAP |
| | IBA1 |
| | MAP2 |
| | Myelin basic protein |
| | NeuN |
| | Neurofilament light |
| | Olig2 |
| | Synaptophysin |
| | TMEM119 |
| GeoMx Mouse NGS Parkinson's Pathology Panel | Alpha-synuclein |
| | ApoA-I |
| | Calbindin |
| | LRRK2 |
| | Park5 |
| | Park7 |
| | Phospho-Alpha-synuclein (S129) |
| | PINK1 |
| | Tyrosine Hydroxylase |
| GeoMx Mouse NGS Core | CD31 |
| | CD45 |
| | GAPDH |
| | GFP |
| | Histone H3 |
| | Ki-67 |
| | Rat IgG2a |
| | Rat IgG2b |
| | Rb IgG |
| | S6 |

**Supplementary Table S4:** A list of Mouse protein targets by module.
