## Supplementary Table 5 for "Ultra High-Plex Spatial Proteogenomic Investigation of Giant Cell Glioblastoma Multiforme Immune Infiltrates Reveals Distinct Protein and RNA Expression Profiles"

| | Cell Line | Tissue Origin |
|---|---|---|
| 1 | COLO201 | LARGE INTESTINE |
| 2 | DAUDI | HAEMATOPOIETIC AND LYMPHOID |
| 3 | H596 | LUNG |
| 4 | HDLM2 | HAEMATOPOIETIC AND LYMPHOID |
| 5 | HEL | HAEMATOPOIETIC AND LYMPHOID |
| 6 | HS578T | BREAST |
| 7 | HUT78 | HAEMATOPOIETIC AND LYMPHOID |
| 8 | MALME3M | SKIN |
| 9 | OPM2 | HAEMATOPOIETIC AND LYMPHOID |
| 10 | THP1 | HAEMATOPOIETIC AND LYMPHOID |
| 11 | U118MG | BRAIN |

**Supplementary Table S5:** The 11-cell pellet array (CPA) used in assay development.
