## Supplementary Table 6 for "Ultra High-Plex Spatial Proteogenomic Investigation of Giant Cell Glioblastoma Multiforme Immune Infiltrates Reveals Distinct Protein and RNA Expression Profiles"

| | Cell Line | Tissue Origin |
|---|---|---|
| 1 | A431 | SKIN |
| 2 | HEK293ICOS | KIDNEY |
| 3 | HEK293PD1 | KIDNEY |
| 4 | HEK293CTLA4 | KIDNEY |
| 5 | HEK293GITR | KIDNEY |
| 6 | HEKLag3 | KIDNEY |
| 7 | HEK293PDL2 | KIDNEY |
| 8 | HEK293PDL1 | KIDNEY |
| 9 | HEK29341BB | KIDNEY |
| 10 | HEK293TIM3 | KIDNEY |
| 11 | CCRF-CEM | PERIPHERAL |
| 12 | 22rv1 | PROSTATE |
| 13 | H596 | LUNG |
| 14 | HCC78 | LUNG |
| 15 | HCT116 | LARGE INTESTINE |
| 16 | HL-60 | PERIPHERAL |
| 17 | HUH7 | HAEMATOPOIETIC AND LYMPHOID |
| 18 | HUT78 | HAEMATOPOIETIC AND LYMPHOID |
| 19 | MDA-MB-468 | BREAST |
| 20 | SK-MEL-5 | SKIN |
| 21 | SK-BR3 | BREAST |
| 22 | RAMOS | HAEMATOPOIETIC AND LYMPHOID |
| 23 | RPMI-8226 | HAEMATOPOIETIC AND LYMPHOID |
| 24 | SU-DHL-6 | PERITONEAL |
| 25 | Ri-1 | HAEMATOPOIETIC AND LYMPHOID |
| 26 | Raji | HAEMATOPOIETIC AND LYMPHOID |
| 27 | OVCAR8 | OVARY |
| 28 | K-562 | HAEMATOPOIETIC AND LYMPHOID |
| 29 | SU-DHL-1 | HAEMATOPOIETIC AND LYMPHOID |
| 30 | SU-DHL-4 | PERITONEAL |
| 31 | SK-MEL-2 | SKIN |
| 32 | WSU-NHL | HAEMATOPOIETIC AND LYMPHOID |
| 33 | THP-1 | PERIPHERAL |
| 34 | SUP-B15 | HAEMATOPOIETIC AND LYMPHOID |
| 35 | U87-MG | BRAIN |
| 36 | U251-MG | BRAIN |
| 37 | NB4 | HAEMATOPOIETIC AND LYMPHOID |
| 38 | SKBR3 PI | BREAST |
| 39 | SH-SY5Y CA | BRAIN |
| 40 | A431 CA | SKIN |
| 41 | NCI-H2228 | LUNG |
| 42 | DBTRG-05MG | BRAIN |
| 43 | SW48 | LARGE INTESTINE |
| 44 | NK-92 | PERIPHERAL |
| 45 | BT-474 | BREAST |

**Supplementary Table S6:** The 45-cell pellet array (CPA) used in assay development.
