## Supplementary Figure 1 for "Ultra High-Plex Spatial Proteogenomic Investigation of Giant Cell Glioblastoma Multiforme Immune Infiltrates Reveals Distinct Protein and RNA Expression Profiles"

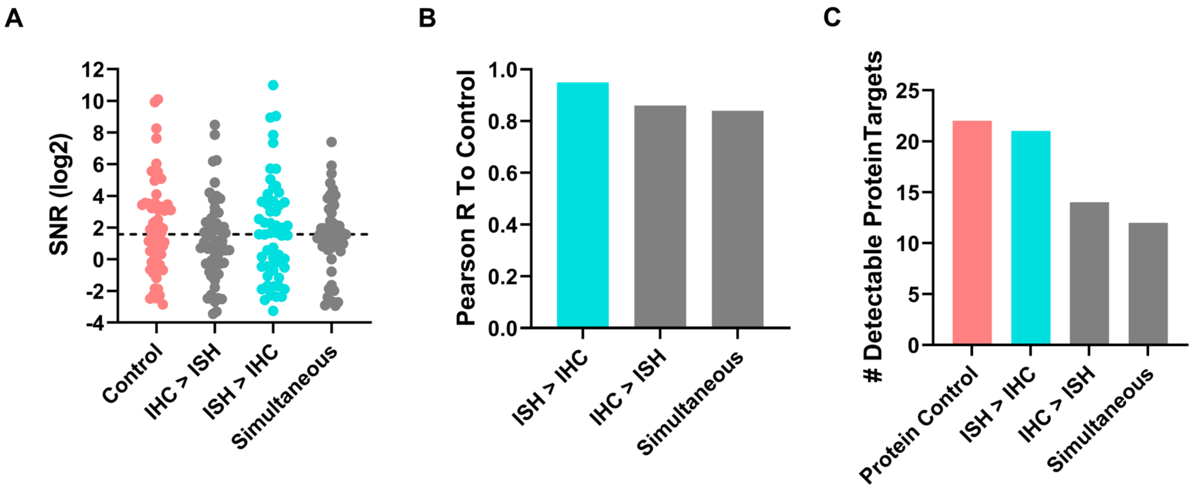


D

| Staining Order | Pearson's Correlation to Protein Control | Decrease in Detectable Targets (%)* |
|---|---|---|
| Mock RNA > Protein (ISH > IHC) | 0.95 | 5 |
| Protein > Mock RNA (IHC > ISH) | 0.86 | 36 |
| Mock RNA:Protein (Simultaneous) | 0.84 | 45 |

**Supplementary Figure S1: Impact of staining order on protein detection.** FFPE cell line, A431CA, was stained with GeoMx Protein assays for nCounter readout and mock RNA probe (Buffer R only). (**A**) Swarm plots showing the distribution of target in relation to SNR. (**B**) Pearson's *R* correlation to the control was calculated for each test condition. (**C**) The number of targets above the detection threshold, SNR $\geq$ 3. (**D**) Summary of the Pearson's *R* correlation and percent decrease in detectable targets for each condition compared to the control.
