## Supplementary Figure 3 for "Ultra High-Plex Spatial Proteogenomic Investigation of Giant Cell Glioblastoma Multiforme Immune Infiltrates Reveals Distinct Protein and RNA Expression Profiles"

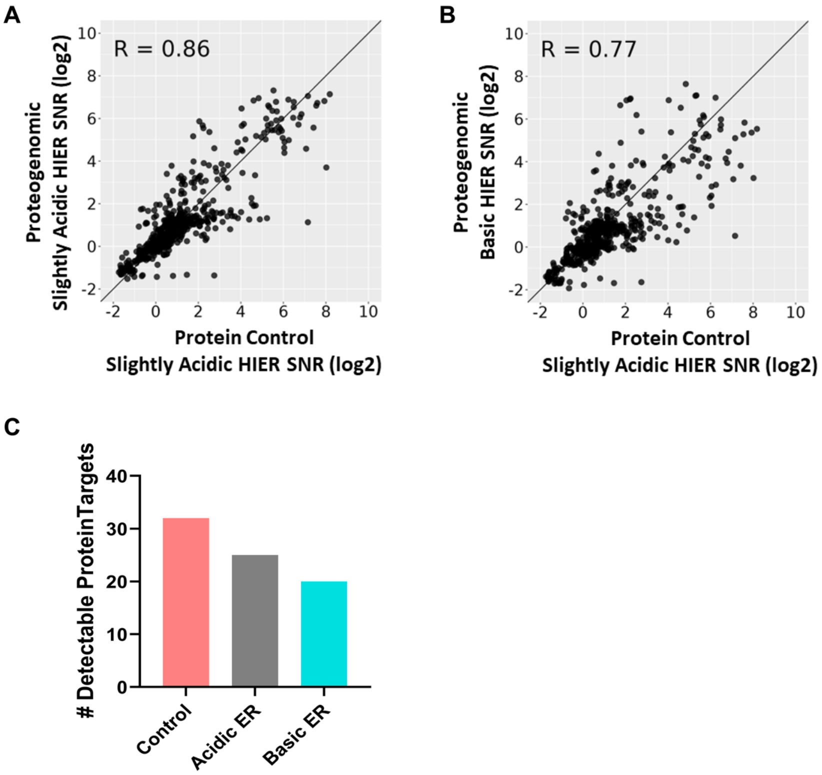


**Supplementary Figure S3: Assessment of epitope retrieval conditions on the performance of spatial proteogenomic assay in relation to protein control.** A 11-core FFPE cell pellet array (CPA) sections, pretreated under basic or slightly acidic conditions and 1 µg/mL proteinase K, were stained with the GeoMx Human CTA and 59-plex GeoMx Human NGS Protein panels. CPAs stained with single-analyte 59-plex GeoMx Human NGS Protein panels under standard assay conditions was used as the protein control. Circular ROIs of 200 µm diameter were selected for detailed molecular profiling with the GeoMx DSP. The signal was averaged across replicate ROIs and the SNR was calculated. Plots represent the correlation of log2-transformed SNR for protein analyte under (**A**) slightly acidic HIER and (**B**) basic HIER. (**C**) The number of protein targets above detection threshold, SNR $\geq$ 3.
