## Supplementary Figure 4 for "Ultra High-Plex Spatial Proteogenomic Investigation of Giant Cell Glioblastoma Multiforme Immune Infiltrates Reveals Distinct Protein and RNA Expression Profiles"

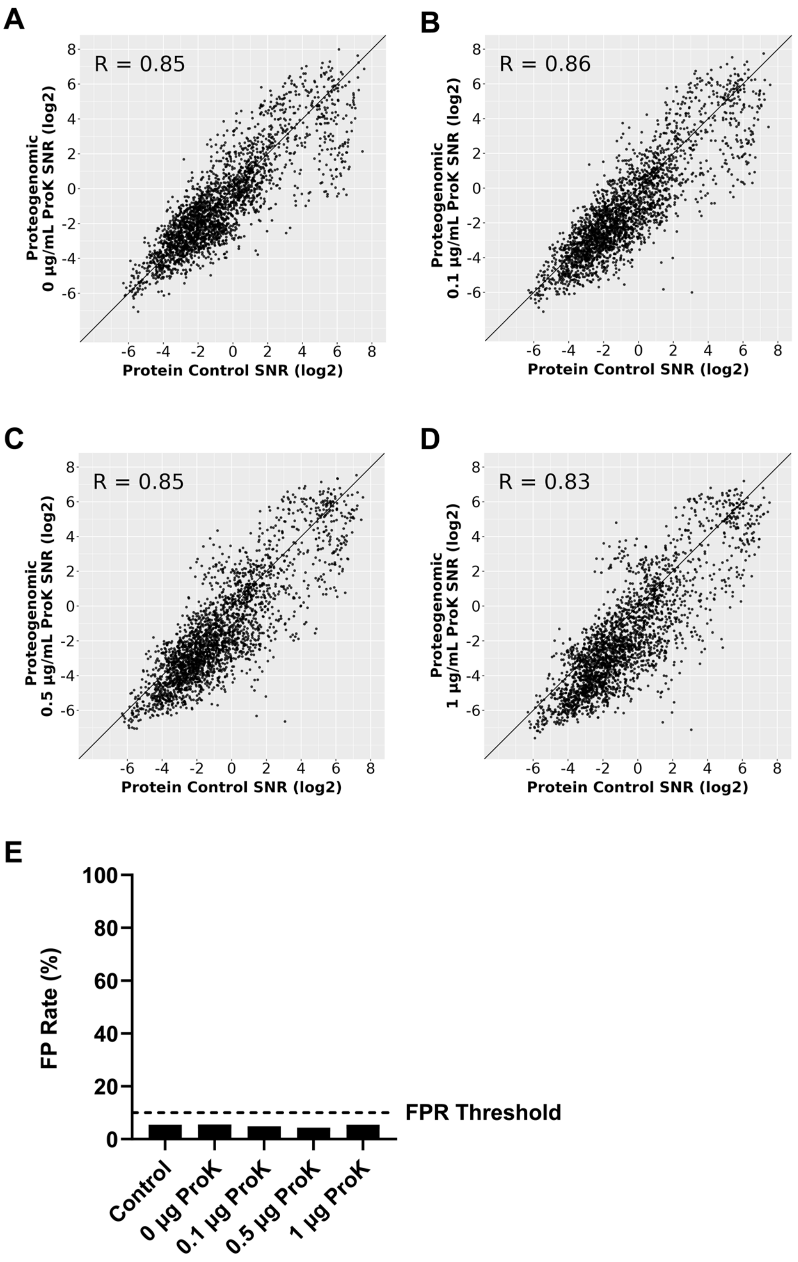


**Supplementary Figure S4: Assessment of varying proteinase K (ProK) on assay performance with respect to protein analyte.** FFPE sections of a 45-cell pellet array was treated under basic HIER followed by varying concentration of ProK. Pretreated slides were stained with GeoMx Human WTA and 59-plex GeoMx Human NGS Protein panels. CPAs stained with single-analyte 59-plex GeoMx Human NGS Protein panels under standard assay conditions was used as the protein control. Circular ROIs of 200 µm diameter were selected for detailed molecular profiling with the GeoMx DSP. The signal was averaged across replicate ROIs and the SNR was calculated. Plots represent the correlation of log2-transformed SNR between the protein control and proteogenomic assay (**A**) 0 µg/mL, (**B**) 0.1 µg/mL, (**C**) 0.5 µg/mL, and (**D**) 1µg/mL ProK. (**E**) Calculated false positive rate (FPR) and of protein targets for each test condition. Dotted line specifies an SNR of 3.
