## Supplementary Figure 5 for "Ultra High-Plex Spatial Proteogenomic Investigation of Giant Cell Glioblastoma Multiforme Immune Infiltrates Reveals Distinct Protein and RNA Expression Profiles"

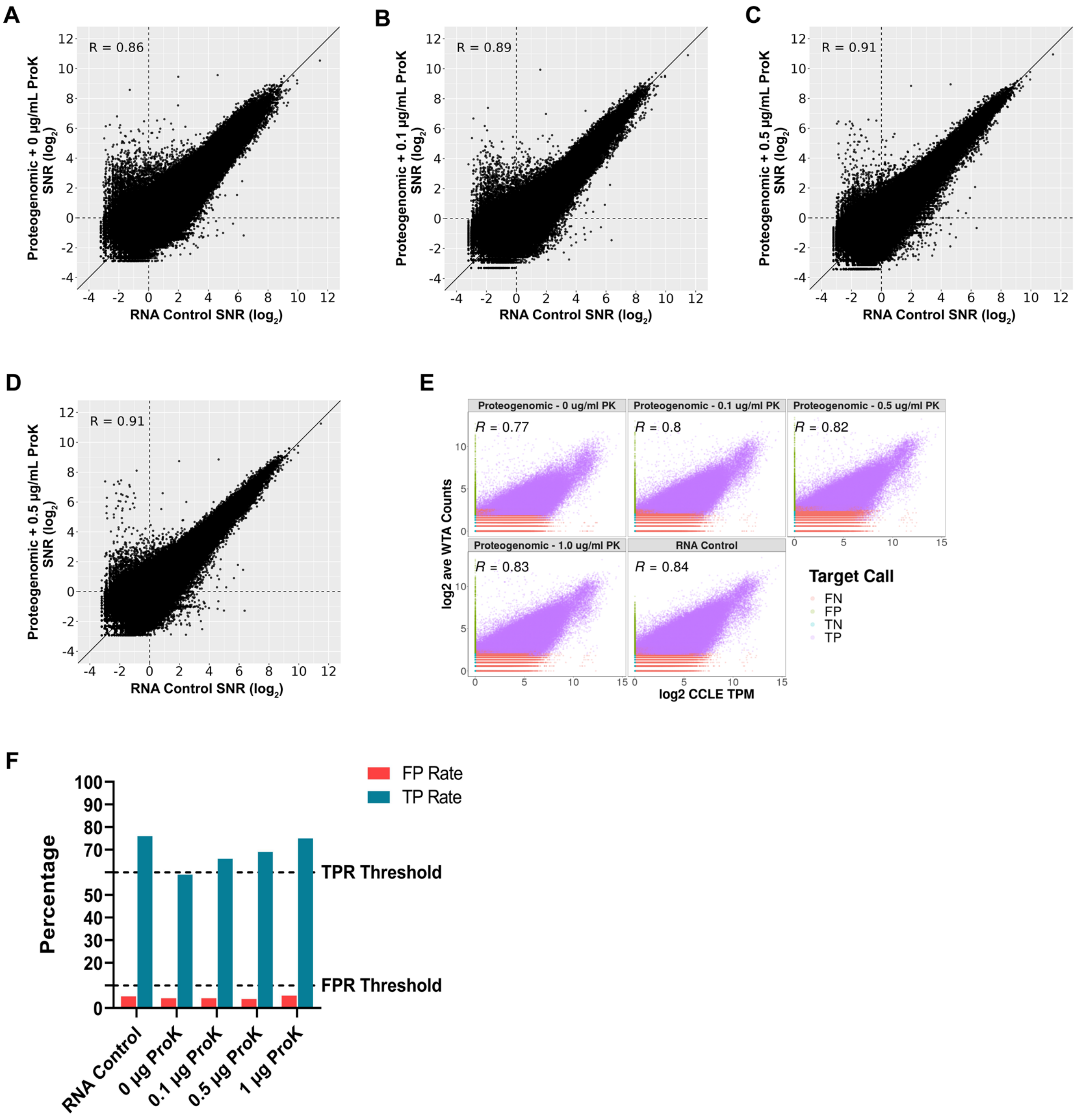


**Supplementary Figure S5: Assessment of varying proteinase K (ProK) on assay performance with respect to RNA analyte.** FFPE sections of a 45-cell pellet array was treated under basic HIER followed by varying concentration of ProK. Pretreated slides were stained with the GeoMx Human WTA and the 59-plex GeoMx Human NGS Protein panels. CPAs stained with the single-analyte GeoMx Human WTA under standard assay conditions was used as the RNA control. Circular ROIs of 200 µm diameter were selected for detailed molecular profiling with the GeoMx DSP. The signal was averaged across replicate ROIs and the SNR was calculated. Plots represent the correlation of log2-transformed SNR between the RNA control and proteogenomic assay (**A**) 0 µg/mL, (**B**) 0.1 µg/mL, (**C**) 0.5 µg/mL, and (**D**) 1 µg/mL ProK. (**E**) Whole slide correlation between the CCLE database and the proteogenomic assay pretreated under varying concentration of ProK and RNA control. (**F**) Calculated false positive rate (FPR) and true positive rate (TPR) for each of the test conditions.
