## Supplementary Figure 6 for "Ultra High-Plex Spatial Proteogenomic Investigation of Giant Cell Glioblastoma Multiforme Immune Infiltrates Reveals Distinct Protein and RNA Expression Profiles"

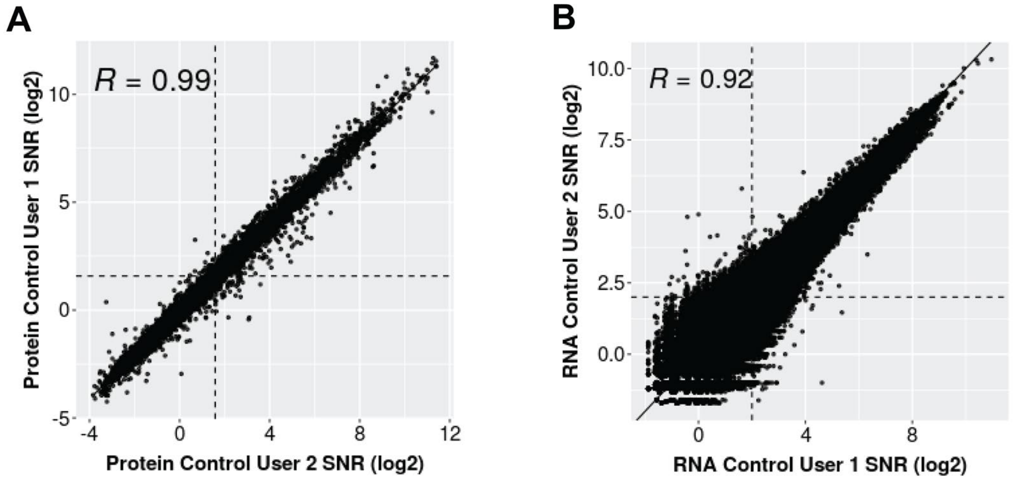


**Supplementary Figure S6: User-to-user and instrument-to-instrument reproducibility for the single analyte** (**A**) protein and (**B**) RNA control.
