## Supplementary Figure 7 for "Ultra High-Plex Spatial Proteogenomic Investigation of Giant Cell Glioblastoma Multiforme Immune Infiltrates Reveals Distinct Protein and RNA Expression Profiles"

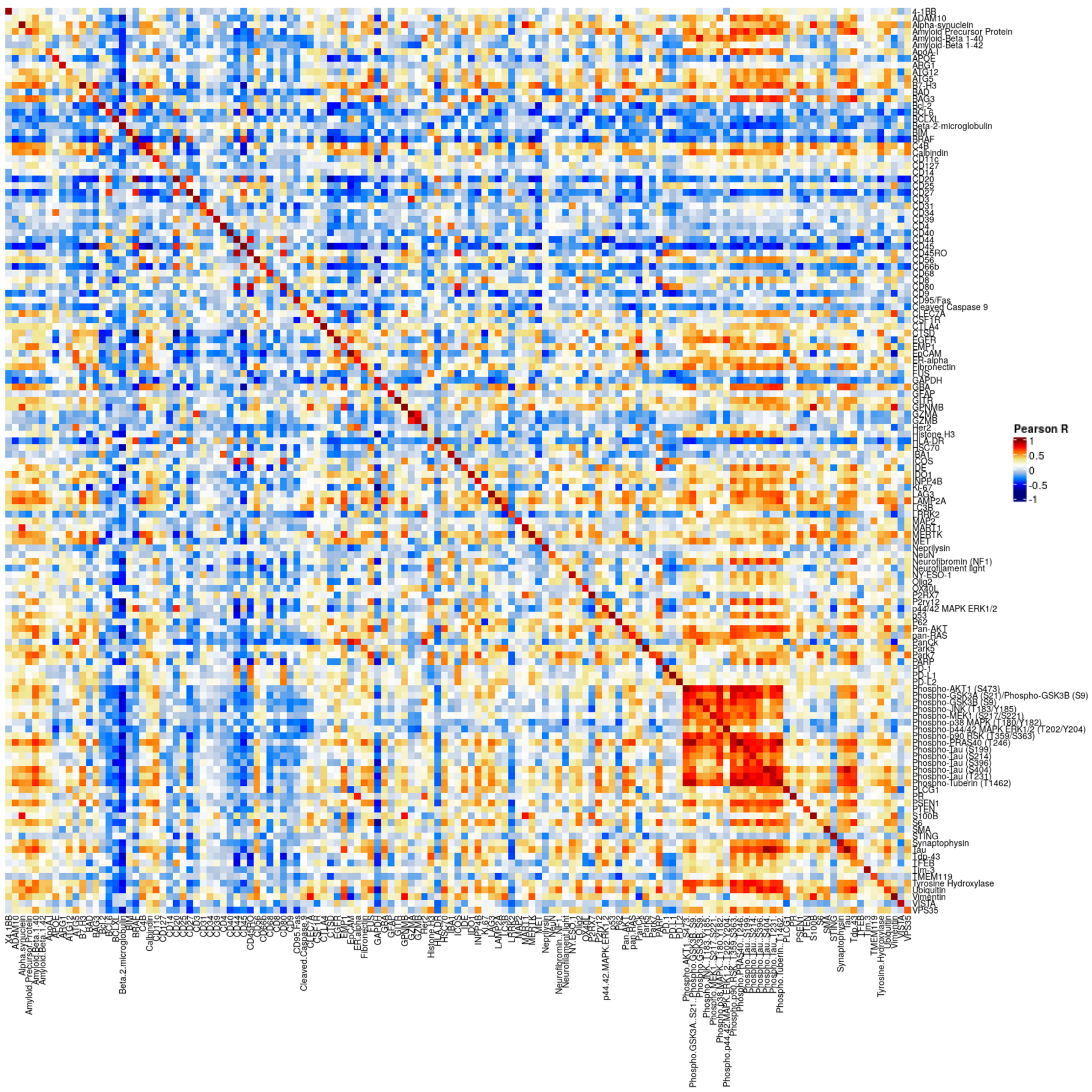


**Supplementary Figure S7: Target to target comparison of Protein Control to proteogenomic protein data.** For protein targets with SNR $\geq$ 3, the Pearson's *R* between each protein target from the Protein Control slide were calculated against all targets in the spatial proteogenomic slide. Heatmap of *R* values are displayed.
