## Supplementary Figure 8 for "Ultra High-Plex Spatial Proteogenomic Investigation of Giant Cell Glioblastoma Multiforme Immune Infiltrates Reveals Distinct Protein and RNA Expression Profiles"

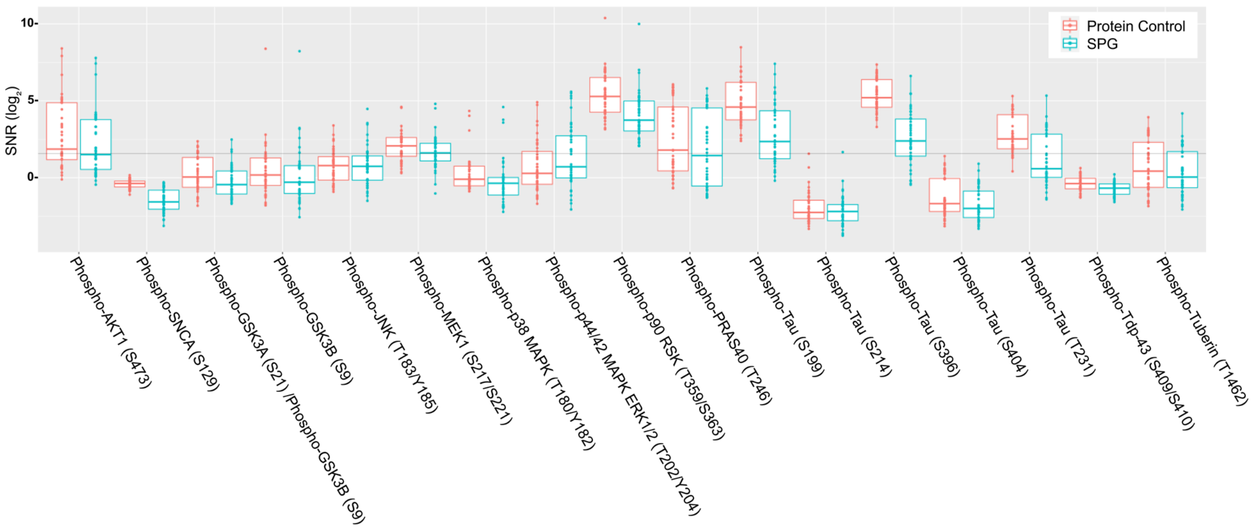


**Supplementary Figure S8: Box plot of the signal-to-noise (SNR) of 17 phospho-specific antibodies under standard and proteogenomic conditions for two separate users.**
