## Supplementary Figure 9 for "Ultra High-Plex Spatial Proteogenomic Investigation of Giant Cell Glioblastoma Multiforme Immune Infiltrates Reveals Distinct Protein and RNA Expression Profiles"

A

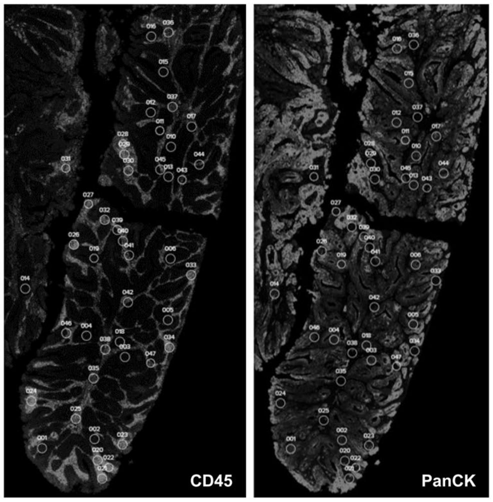


B

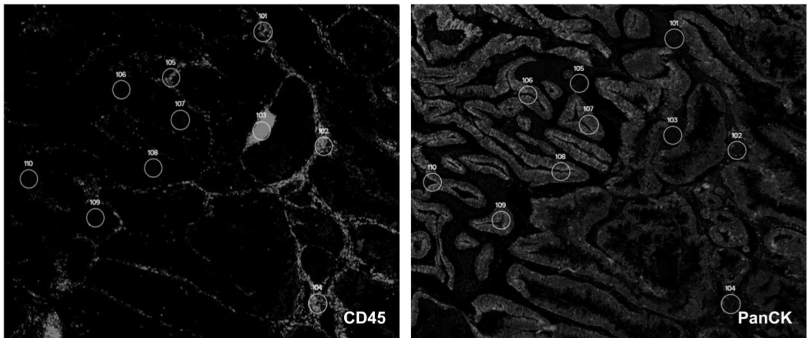


**Supplementary Figure S9: Representative images of CRC sample used in the assessment of spatial proteogenomic data quality.** FFPE colorectal cancer (CRC) sections were stained with the GeoMx NGS Human Protein modules (147-plex), WTA, and antibodies against CD45 (Immune; left panel) and PanCK (Tumor; right panel).
