## Supplementary Figure 10 for "Ultra High-Plex Spatial Proteogenomic Investigation of Giant Cell Glioblastoma Multiforme Immune Infiltrates Reveals Distinct Protein and RNA Expression Profiles"

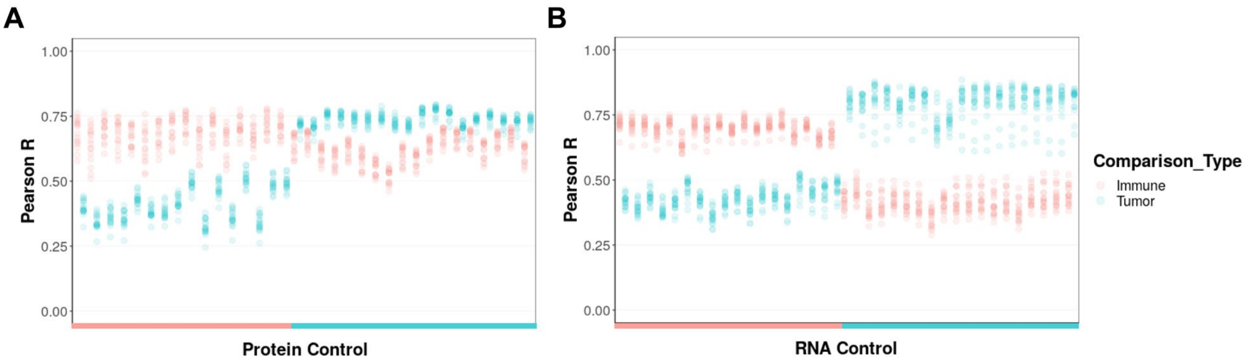


**Supplementary Figure S10: ROI-to-ROI comparison of the proteogenomic data to the single analyte controls.** CRC FFPE sections were stained with 147-plex GeoMx NGS human Protein modules, WTA, and antibodies against PanCK (Tumor) and CD45 (Immune). The Pearson's *R* was calculated between each ROI from the proteogenomic assay against all ROIs in the single analyte (**A**) protein and (**B**) RNA controls. ROIs are colored according to region (immune or tumor).
