## Supplementary Figure 11 for "Ultra High-Plex Spatial Proteogenomic Investigation of Giant Cell Glioblastoma Multiforme Immune Infiltrates Reveals Distinct Protein and RNA Expression Profiles"

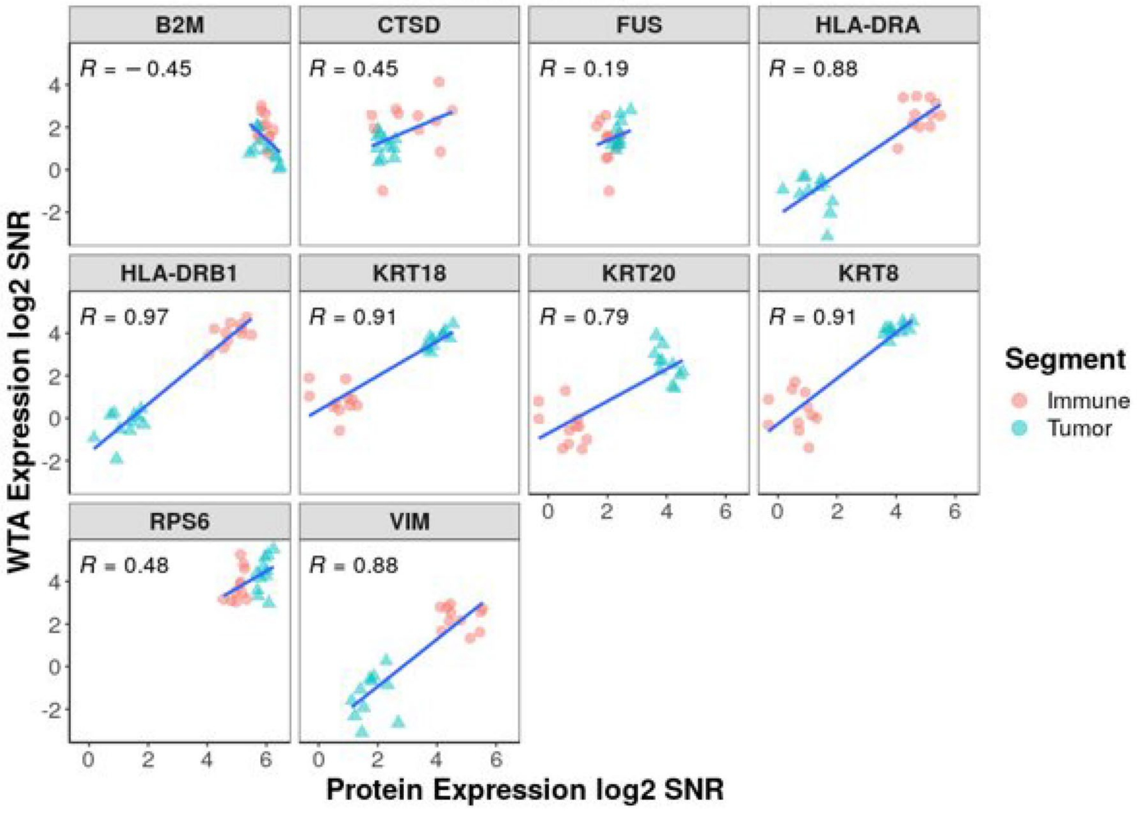


**Supplementary Figure S11: Concordance between matching protein and RNA targets.** For protein targets with SNR ≥ 3 and the respective RNA target with SNR ≥ 4, a pairwise scatterplot was generated to visualize the concordance between respective analytes. Pearson's *R* calculations are shown in each plot.
