## Supplementary Figure 12 for "Ultra High-Plex Spatial Proteogenomic Investigation of Giant Cell Glioblastoma Multiforme Immune Infiltrates Reveals Distinct Protein and RNA Expression Profiles"

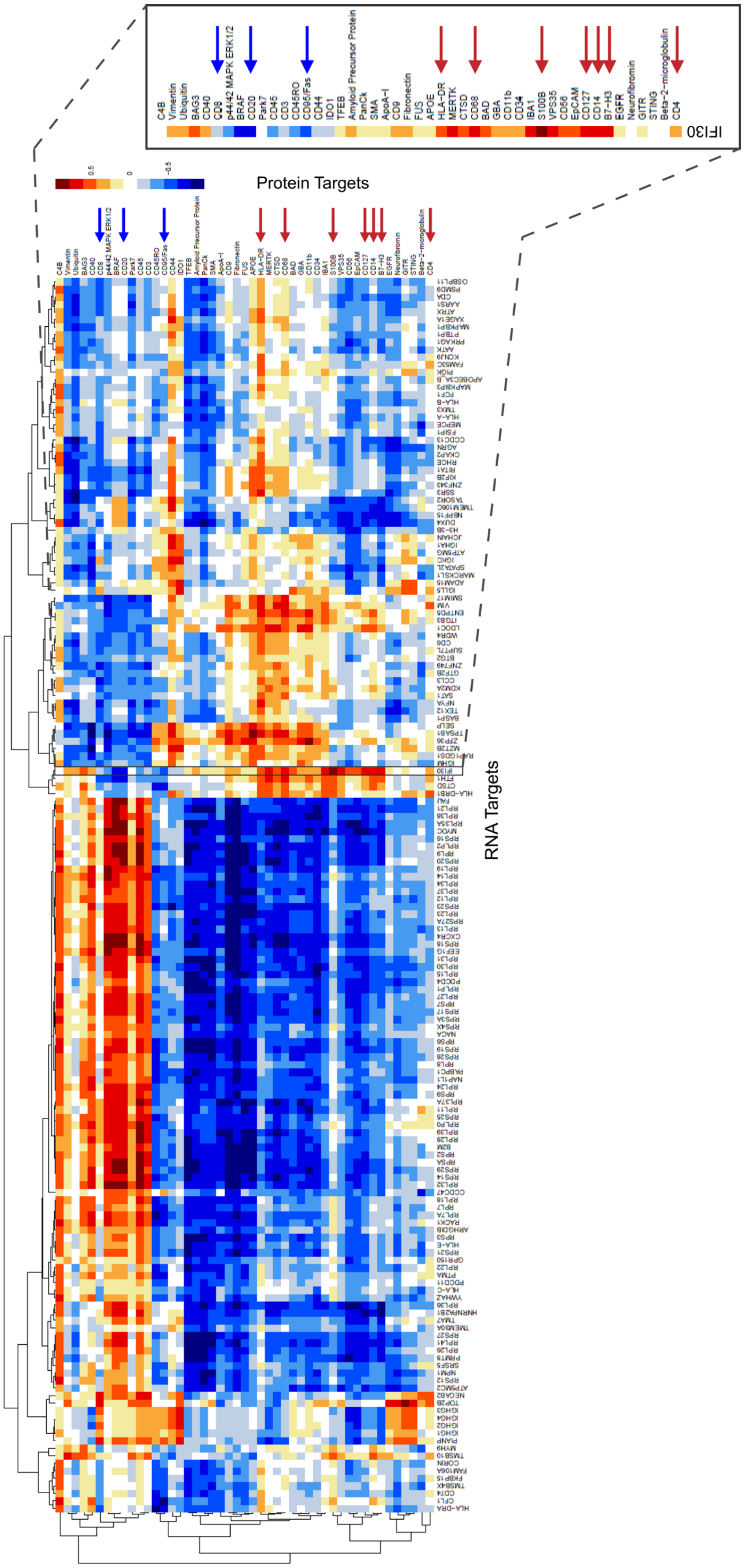
Protein Targets
RNA Targets
IFI30

**Supplementary Figure S12: Concordance between Proteogenomic RNA and Protein targets above background in immune segments of colorectal cancer (CRC).** FFPE section of CRC stained with 147-plex GeoMx NGS Protein modules, WTA, and antibodies against PanCK (Tumor) and CD45 (Immune) using the proteogenomic workflow. Tumor and immune segments were selected based on PanCK and CD45 immunofluorescence, respectively. Pearson's *R* was calculated between each detected protein target (SNR $\geq$ 3) and RNA targets (SNR $\geq$ 4) within the immune segment.
