## Supplementary Figure 13 for "Ultra High-Plex Spatial Proteogenomic Investigation of Giant Cell Glioblastoma Multiforme Immune Infiltrates Reveals Distinct Protein and RNA Expression Profiles"

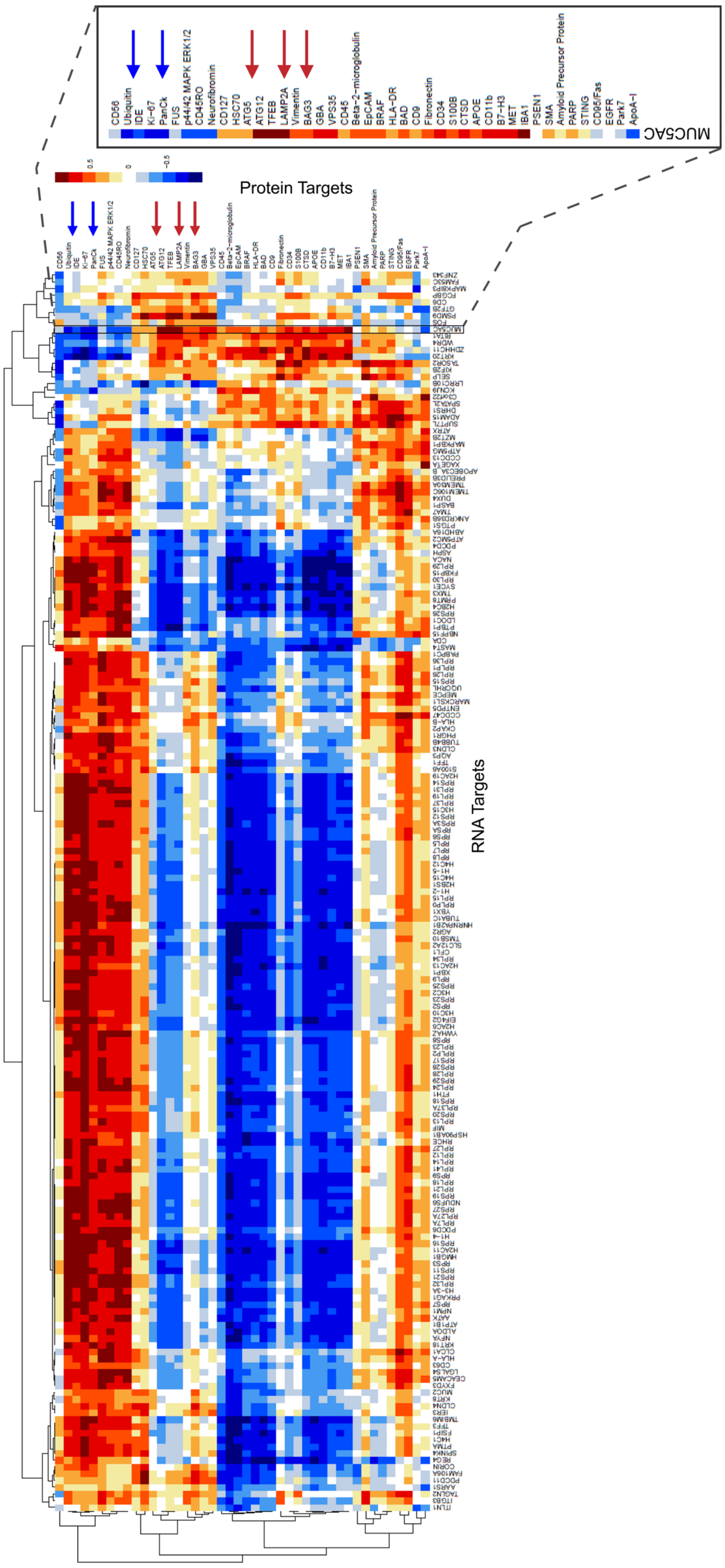

CD56
Ubiquitin
IDE
Ki-67
PanCk
FUS
p44/42 MAPK ERK1/2
CD45RO
Neurofibromin
CD127
HSC70
ATG5
ATG12
TFEB
LAMP2A
Vimentin
BAG3
GBA
VPS35
CD45
Beta-2-microglobulin
EpCAM
BRAF
HLA-DR
BAD
CD9
Fibronectin
CD34
S100B
CTSD
APOE
CD11b
B7-H3
MET
IBA1
PSEN1
SMA
Amyloid Precursor Protein
PARP
STING
CD95/Fas
EGFR
Park7
ApoA-I
MUC5AC
0.5
0
-0.5
Protein Targets
RNA Targets

**Supplementary Figure S13: Concordance between Proteogenomic RNA and Protein targets above background in tumor segments of colorectal cancer (CRC).** FFPE section of CRC stained with 147-plex GeoMx NGS Protein modules, WTA, and antibodies against PanCK (Tumor) and CD45 (Immune) using the proteogenomic workflow. Tumor and immune segments were selected based on PanCK and CD45 immunofluorescence, respectively. Pearson's *R* was calculated between each detected protein target (SNR $\geq$ 3) and RNA targets (SNR $\geq$ 4) within the tumor segment.
