## Supplementary Figure 14 for "Ultra High-Plex Spatial Proteogenomic Investigation of Giant Cell Glioblastoma Multiforme Immune Infiltrates Reveals Distinct Protein and RNA Expression Profiles"

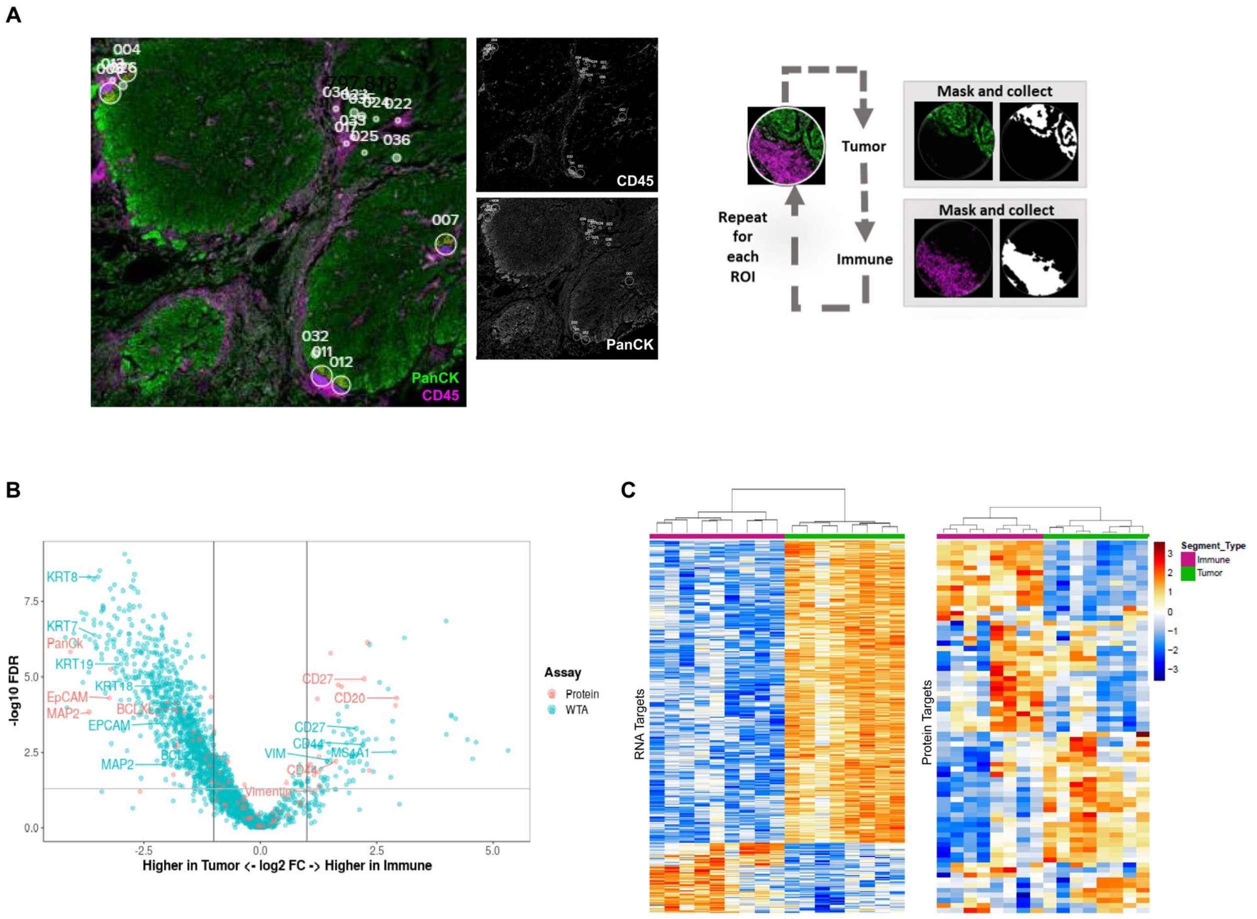


**Supplementary Figure S14: Assessment of spatial proteogenomic performance on human NSCLC.** NSCLC FFPE sections were stained with 147-plex GeoMx NGS human Protein modules, WTA, and antibodies against PanCK (Tumor) and CD45 (Immune). (**A**) Multiplexed protein and RNA characterization of NSCLC sample with representative colored and gray scaled images highlighting the segmentation of 300 µm circular ROIs into tumor (PanCK$^{+}$) and immune (CD45$^{+}$) enriched regions. Segments illuminated in white were collected, black regions were not. Protein and RNA counts were SNR transformed and protein targets with SNR $\geq$ 3 and WTA RNA targets with SNR $\geq$ 4 were used in the analysis. (**B**) Combined volcano plot of protein and RNA expression in NSCLC. All immune segments were compared to all tumor segments for protein and RNA targets above background. A subset of differentially expressed genes are labeled with colors matching their analyte. (**C**) Unsupervised hierarchical clustering of detected RNA (left) and protein (right) targets for NSCLC.
