## Supplementary Figure 15 for "Ultra High-Plex Spatial Proteogenomic Investigation of Giant Cell Glioblastoma Multiforme Immune Infiltrates Reveals Distinct Protein and RNA Expression Profiles"

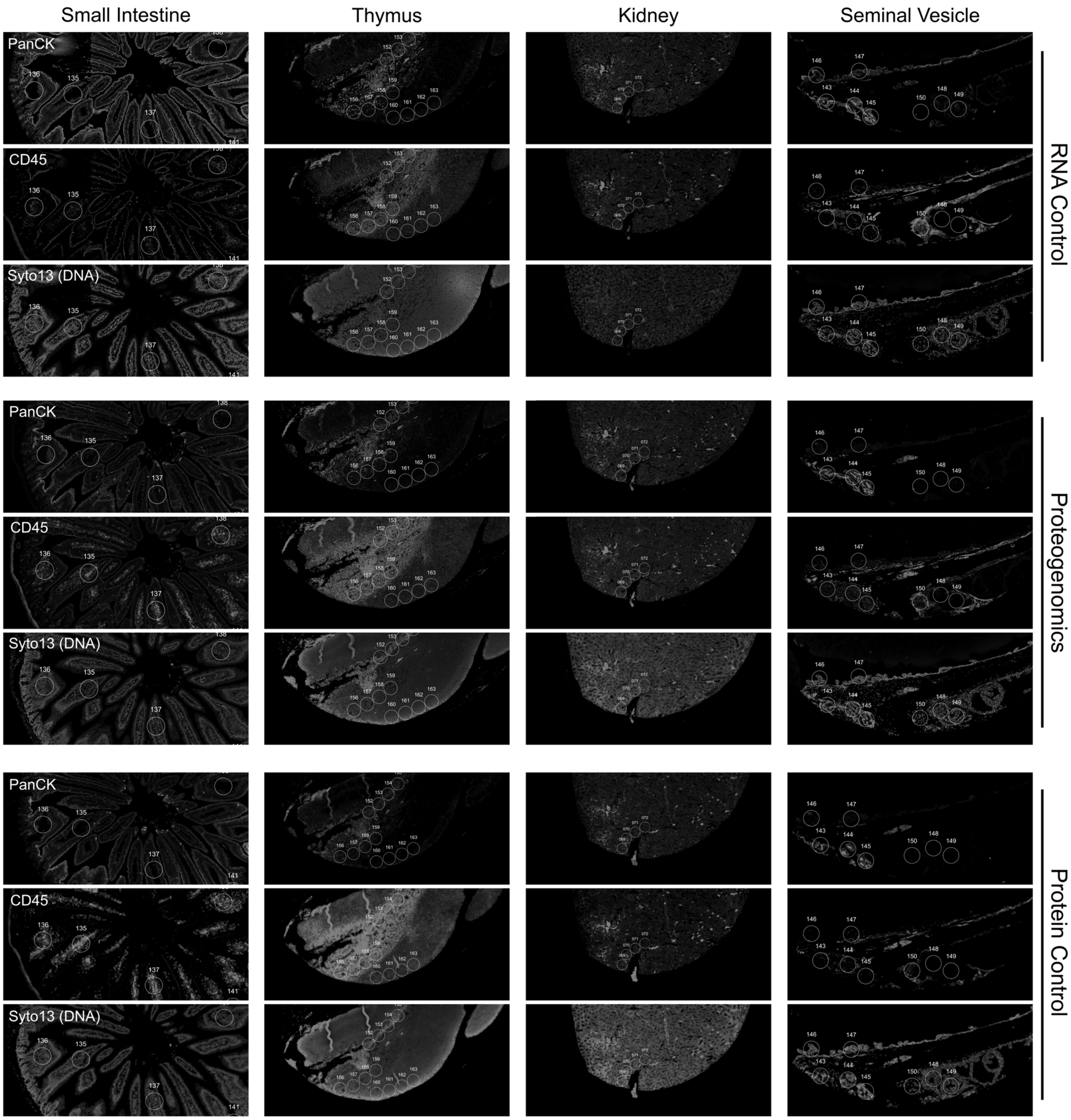


**Supplementary Figure S15: Representative grey scale images of used in the assessment of spatial proteogenomic data quality across mouse tissue types.** FFPE sections were stained with the GeoMx NGS Mouse Protein modules (137-plex), GeoMx Mm WTA, and antibodies against CD45 (Immune), PanCK (Tumor) and Syto13 (nuclear, DNA).
