## Supplementary Figure 16 for "Ultra High-Plex Spatial Proteogenomic Investigation of Giant Cell Glioblastoma Multiforme Immune Infiltrates Reveals Distinct Protein and RNA Expression Profiles"

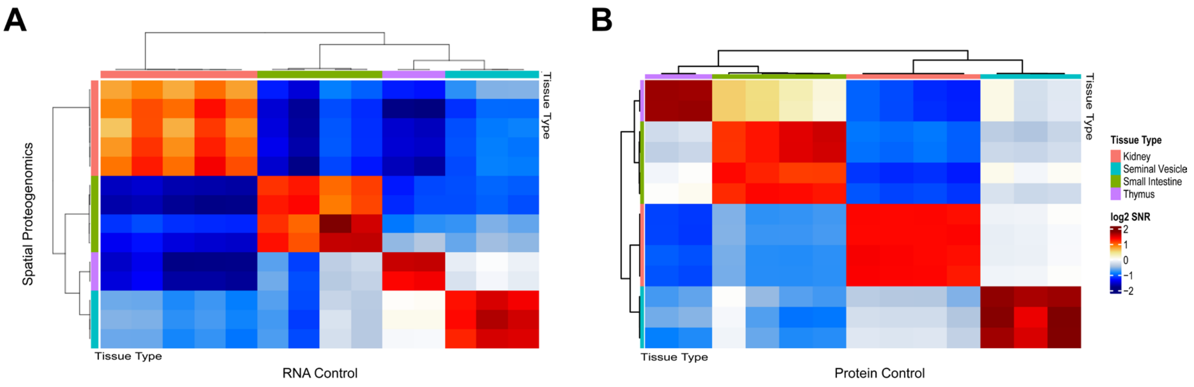


**Supplementary Figure S16: ROI-to-ROI comparison of the proteogenomic data to the single analyte controls.** Mouse FFPE sections were stained with 137-plex GeoMx NGS Mouse Protein modules and GeoMx Mm WTA. The Pearson's *R* was calculated between each ROI from the proteogenomic assay against all ROIs in the single analyte (**A**) RNA and (**B**) protein controls.
