## Supplementary Figure 17 for "Ultra High-Plex Spatial Proteogenomic Investigation of Giant Cell Glioblastoma Multiforme Immune Infiltrates Reveals Distinct Protein and RNA Expression Profiles"

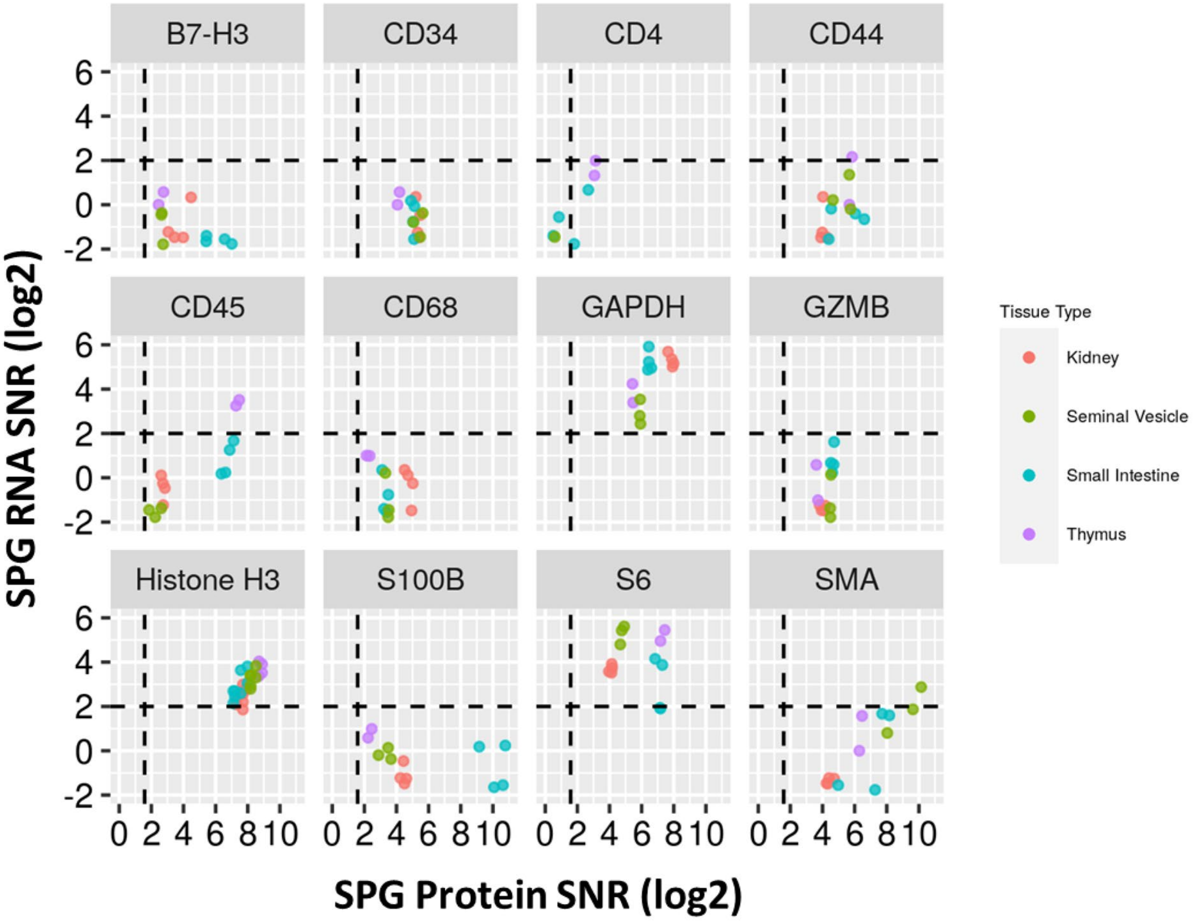


**Supplementary Figure S17: Expression levels of select matching RNA and protein targets in several mouse tissue types.** For protein targets with SNR ≥ 3 and the respective RNA target with SNR ≥ 4, a pairwise scatter plot was generated to visualize the concordance between respective analytes.
