## Supplementary Figure 18 for "Ultra High-Plex Spatial Proteogenomic Investigation of Giant Cell Glioblastoma Multiforme Immune Infiltrates Reveals Distinct Protein and RNA Expression Profiles"

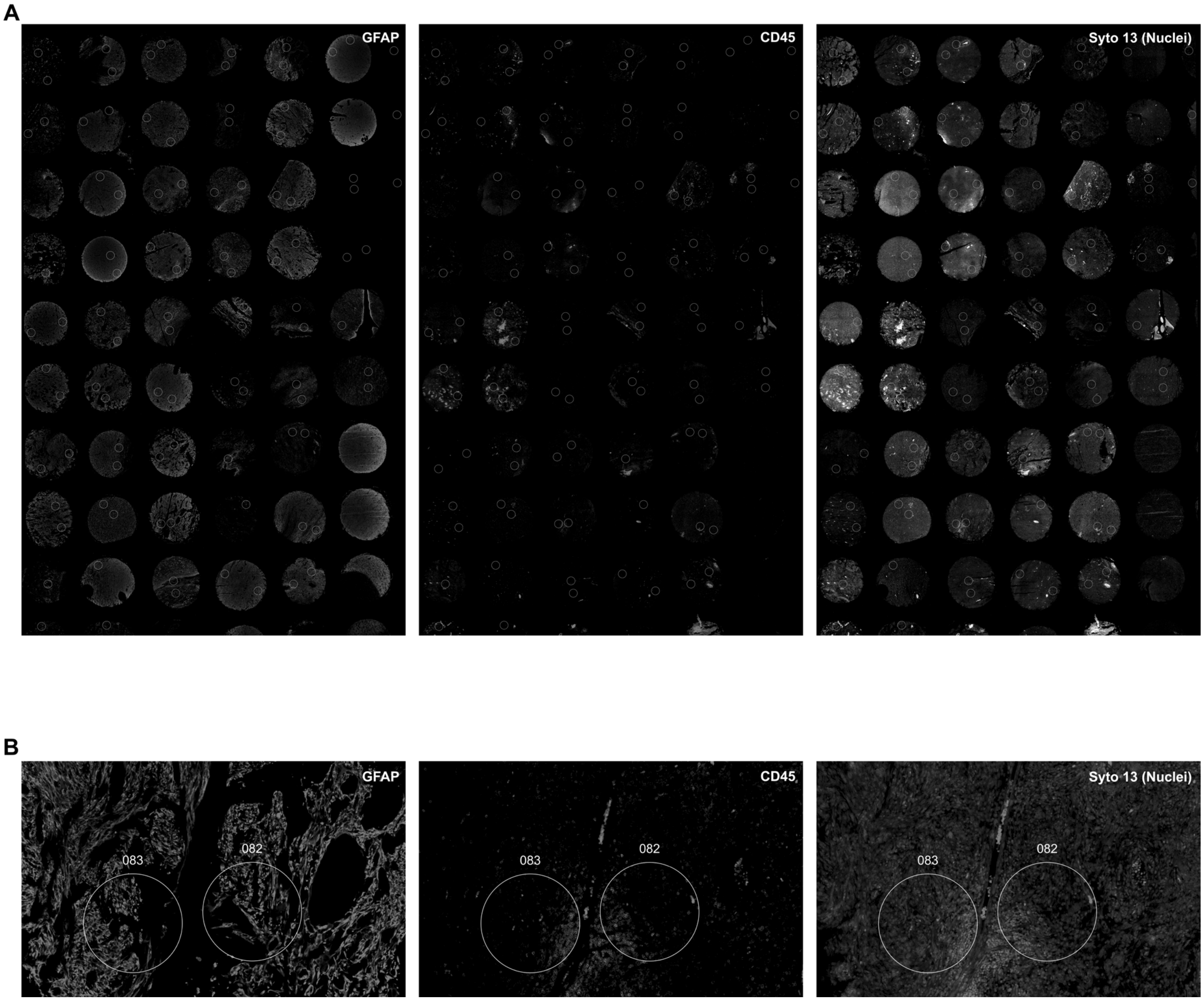


**Supplementary Figure S18: Representative images used in the assessment of spatial proteogenomic data quality on Glioblastoma multiforme Grade 4.** (A) Human brain glioblastoma tissue array was stained with the GeoMx NGS Human Protein modules (147-plex), WTA, and antibodies against GFAP (Astrocyte/Tumor; left panel), CD45 (Immune; middle panel) and Syto13 (Nuclei; right panel). (B) ROI were segmented into GFAP$^+$ (left panel), CD45$^+$ (middle panel) or Syto13 (right panel).
