## Supplementary Figure 19 for "Ultra High-Plex Spatial Proteogenomic Investigation of Giant Cell Glioblastoma Multiforme Immune Infiltrates Reveals Distinct Protein and RNA Expression Profiles"

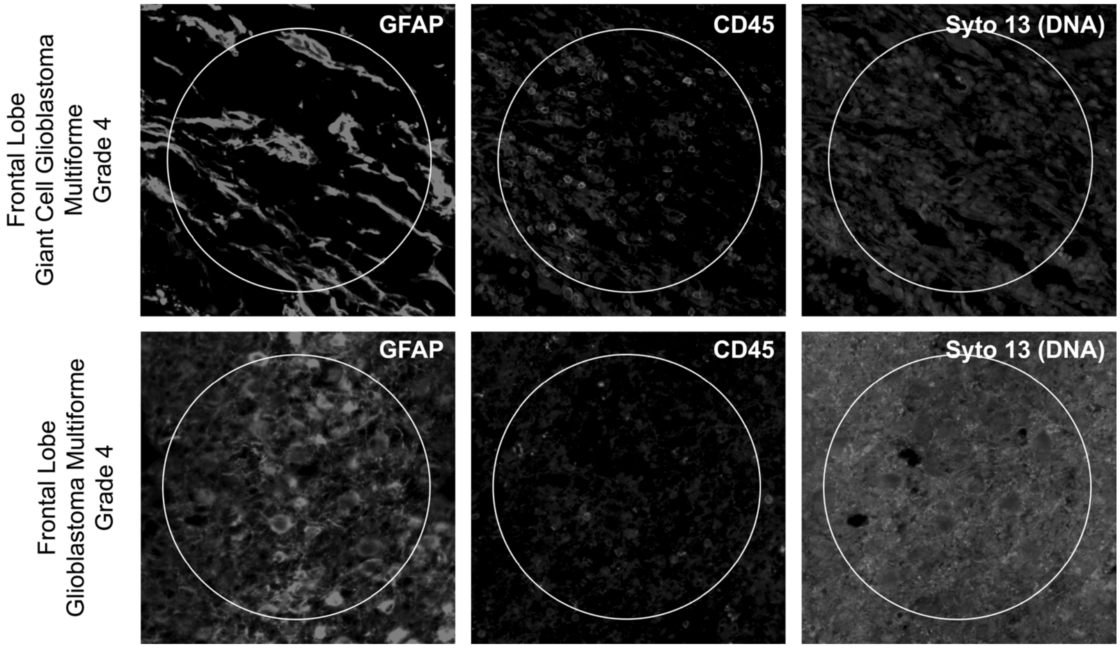


**Supplementary Figure S19: Representative images used in the assessment of spatial proteogenomic data quality on gcGBM and GBM.** ROIs for gcGBM (top panel) and GBM (bottom panel) were segmented into GFAP$^+$ (Astrocyte/Tumor; left panel), CD45$^+$ (Immune; middle panel) and Syto13 (Nuclear, DNA; right panel).
