## Supplementary Figure 20 for "Ultra High-Plex Spatial Proteogenomic Investigation of Giant Cell Glioblastoma Multiforme Immune Infiltrates Reveals Distinct Protein and RNA Expression Profiles"

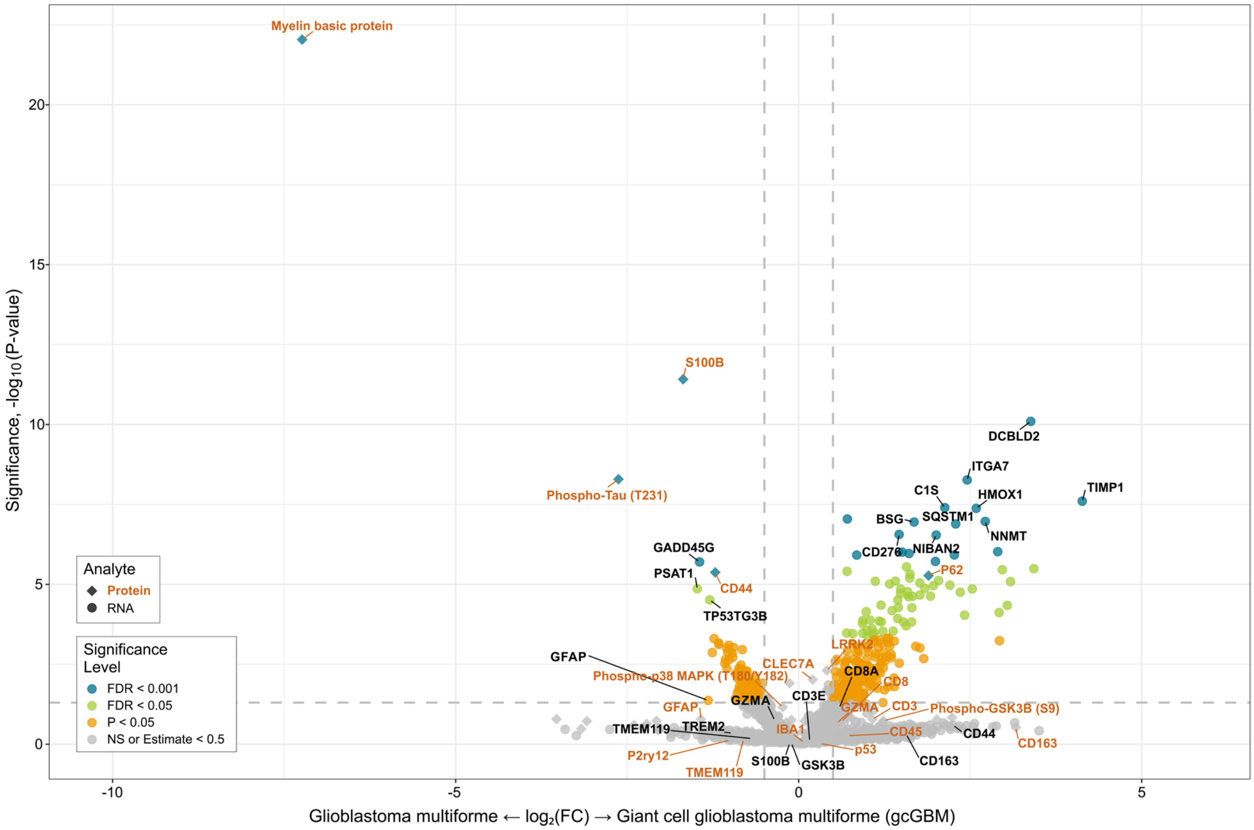


**Supplementary Figure S20: Combined volcano plot resulting from the differential expression analysis between GBM and gcGBM for all AOIs within the left frontal lobe for RNA (●) and protein (♦) analytes.** Targets with significantly differential expression are highlighted either orange (P-value < 0.05), green (FDR < 0.05) or blue (FDR < 0.01); whereas targets in grey show no significant difference in expression. Genes with significantly differential expression are highlighted either orange (P-value < 0.05), green (FDR < 0.05) or blue (FDR <0.01); whereas genes in grey show no significant difference in expression. A subset of differentially expressed genes are labeled.
