## Supplementary Figure 21 for "Ultra High-Plex Spatial Proteogenomic Investigation of Giant Cell Glioblastoma Multiforme Immune Infiltrates Reveals Distinct Protein and RNA Expression Profiles"

A

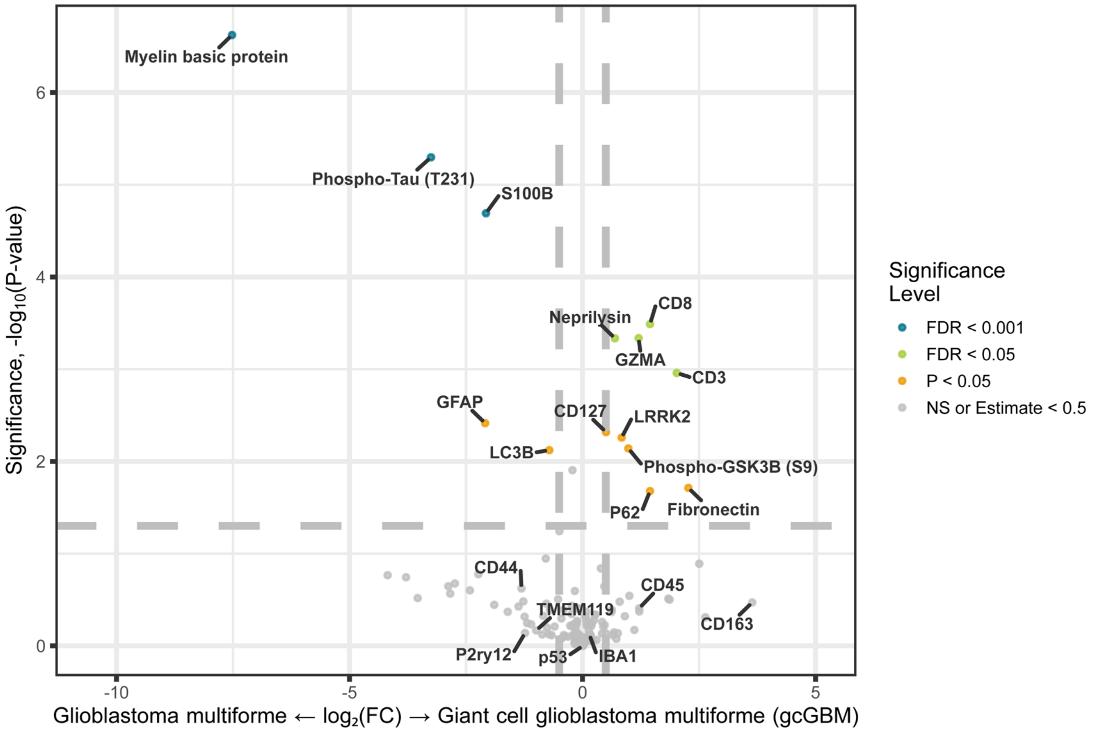
Myelin basic protein
Phospho-Tau (T231)
S100B
CD8
Neprilysin
GZMA
CD3
GFAP
CD127
LRRK2
LC3B
Phospho-GSK3B (S9)
P62
Fibronectin
CD44
CD45
TMEM119
CD163
P2ry12
p53
IBA1
Significance, -log10(P-value)
Glioblastoma multiforme ← log2(FC) → Giant cell glioblastoma multiforme (gcGBM)
Significance Level
FDR < 0.001
FDR < 0.05
P < 0.05
NS or Estimate < 0.5

B

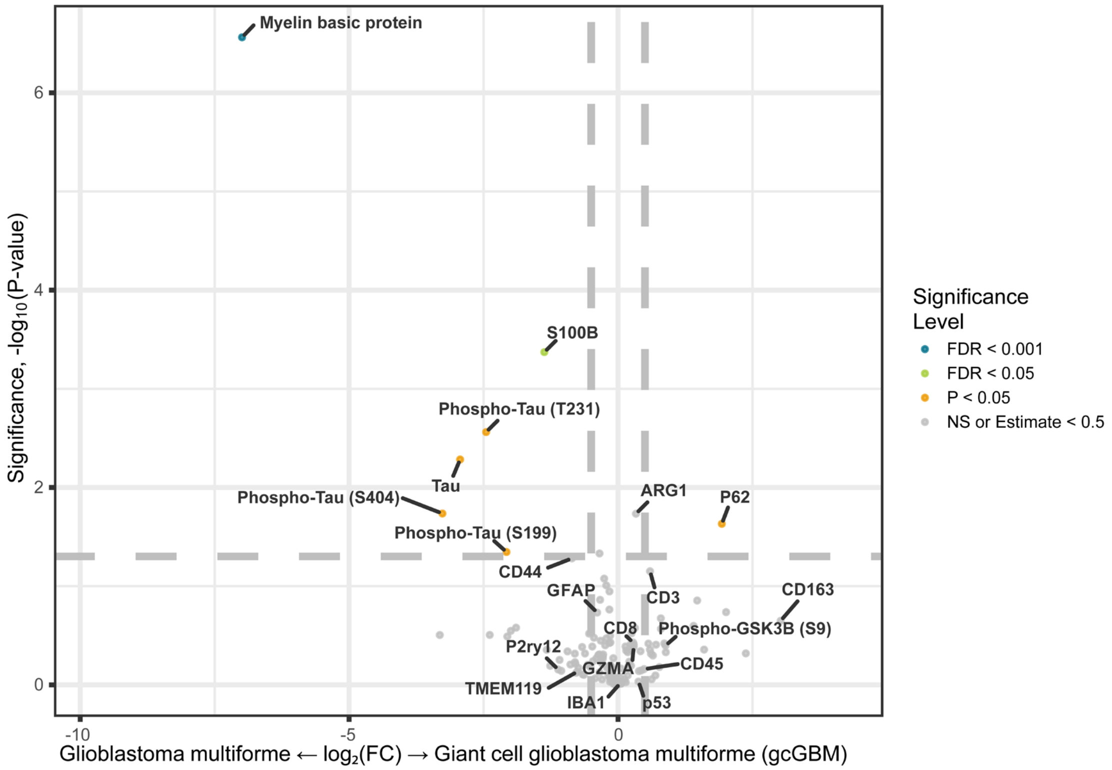
Myelin basic protein
S100B
Phospho-Tau (T231)
Tau
Phospho-Tau (S404)
ARG1
P62
Phospho-Tau (S199)
CD44
GFAP
CD3
CD163
CD8
Phospho-GSK3B (S9)
P2ry12
GZMA
CD45
TMEM119
IBA1
p53
Significance, -log10(P-value)
Glioblastoma multiforme ← log2(FC) → Giant cell glioblastoma multiforme (gcGBM)
Significance Level
FDR < 0.001
FDR < 0.05
P < 0.05
NS or Estimate < 0.5

**Supplementary Figure S21: Differential protein expression analysis between GBM and gcGBM.** Volcano showing the differential protein expression profiles for (**A**) CD45- and (**B**) GFAP-enriched segments within the left frontal lobe. Proteins with significantly differential expression are highlighted either orange (*P*-value < 0.05), green (FDR < 0.05) or blue (FDR < 0.01), whereas proteins in grey show no significant difference in expression. A subset of differentially expressed proteins are labeled.
